# Phables: from fragmented assemblies to high-quality bacteriophage genomes

**DOI:** 10.1101/2023.04.04.535632

**Authors:** Vijini Mallawaarachchi, Michael J. Roach, Przemyslaw Decewicz, Bhavya Papudeshi, Sarah K. Giles, Susanna R. Grigson, George Bouras, Ryan D. Hesse, Laura K. Inglis, Abbey L. K. Hutton, Elizabeth A. Dinsdale, Robert A. Edwards

## Abstract

**Motivation:** Microbial communities influence both human health and different environments. Viruses infecting bacteria, known as bacteriophages or phages, play a key role in modulating bacterial communities within environments. High-quality phage genome sequences are essential for advancing our understanding of phage biology, enabling comparative genomics studies, and developing phage-based diagnostic tools. Most available viral identification tools consider individual sequences to determine whether they are of viral origin. As a result of the challenges in viral assembly, fragmentation of genomes can occur, leading to the need for new approaches in viral identification. Therefore, the identification and characterisation of novel phages remain a challenge.

**Results:** We introduce Phables, a new computational method to resolve phage genomes from fragmented viral metagenome assemblies. Phables identifies phage-like components in the assembly graph, models each component as a flow network, and uses graph algorithms and flow decomposition techniques to identify genomic paths. Experimental results of viral metagenomic samples obtained from different environments show that Phables recovers on average over 49% more high-quality phage genomes compared to existing viral identification tools. Furthermore, Phables can resolve variant phage genomes with over 99% average nucleotide identity, a distinction that existing tools are unable to make.

**Availability and Implementation:** Phables is available on GitHub at https://github.com/Vini2/phables.

## Introduction

Bacteriophages (hereafter *phages*) are viruses that infect bacteria, which influence microbial ecology and help modulate microbial communities (Edwards and Rohwer 2005; Rodriguez-Valera et al. 2009). Phages are considered the most abundant biological entity on earth, totalling an estimated 10^31^ particles (Comeau et al. 2008). Since their discovery by Frederick Twort in 1915 (Twort 1915), phages have been isolated from many diverse environments (Keen 2015). When sequencing technologies were first developed, phage genomes were the first to be sequenced due to their relatively small genome size (Sanger et al. 1977). With the advent of second-generation sequencing technologies, the first metagenomic samples to be sequenced were phages (Breitbart et al. 2002). The availability of advanced sequencing technologies has facilitated the investigation of the effect of phages on the functions of microbial communities, especially in the human body’s niche areas. For example, phages residing in the human gut have a strong influence on human health (Łusiak-Szelachowska et al. 2017) and impact gastrointestinal diseases such as inflammatory bowel disease (IBD) (Norman et al. 2015). To date, our understanding of the diversity of phages is limited, as most have not been cultured due to the inherent difficulty of recovering phages from their natural environments. Although countless millions of phage species are thought to exist, only 25,936 complete phage genomes have been sequenced according to the INfrastructure for a PHAge REference Database (INPHARED) (Cook et al. 2021) (as of the July 2023 update).

Metagenomics has enabled the application of modern sequencing techniques for the culture-independent study of microbial communities (Hugenholtz, Goebel, and Pace 1998). Metagenomic sequencing provides a multitude of sequencing reads from the genetic material in environmental samples that are composed of a mixture of prokaryotic, eukaryotic, and viral species. Metagenomic analysis pipelines start by assembling sequencing reads from metagenomic samples into longer contiguous sequences that are used in downstream analyses. Most metagenome assemblers (D. Li et al. 2015; Nurk et al. 2017; Namiki et al. 2012; Peng et al. 2011) use de Bruijn graphs (P. A. Pevzner, Tang, and Waterman 2001) as the primary data structure where they break sequencing reads into smaller pieces of length k (known as ‘*k*-mers’) and represent *k*-mers as vertices and edges as overlaps of length *k-1*. After performing several simplification steps, the final *assembly graph* represents sequences as vertices and connection information between these sequences as edges (Nurk et al. 2017; V. Mallawaarachchi, Wickramarachchi, and Lin 2020). Non-branching paths in the assembly graph (paths where all vertices have an in-degree and out-degree of one, except for the first and last vertices) are referred to as *unitigs* (Kececioglu and Myers 1995). Unitigs are entirely consistent with the read data and belong to the final genome(s). Assemblers condense unitigs into individual vertices and resolve longer optimised paths through the branches into contiguous sequences known as *contigs* (Bankevich et al. 2012). As the contextual and contiguity information of reads is lost in de Bruijn graphs, mutations in metagenomes with high strain diversity appear as “*bubbles*” in the assembly graph where a vertex has multiple outgoing edges (branches) which eventually converge as incoming edges into another vertex (P. A. Pevzner, Tang, and Waterman 2001; Pavel A. Pevzner, Tang, and Tesler 2004). Assemblers consider these bubbles as errors and consider one path of the bubble corresponding to the dominant strain (Bankevich et al. 2012) or terminate contigs prematurely (D. Li et al. 2015). Moreover, most metagenome assemblers are designed and optimised for bacterial genomes and fail to recover viral populations with low coverage and genomic repeats (Roux et al. 2017; Sutton et al. 2019). However, previous studies have shown that contigs that are connected to each other are more likely to belong to the same genome (V. Mallawaarachchi, Wickramarachchi, and Lin 2020; V. G. Mallawaarachchi, Wickramarachchi, and Lin 2020, 2021). Hence, the assembly graph retains important connectivity and neighbourhood information within fragmented assemblies. This concept has been successfully applied to develop tools such as GraphMB (Lamurias et al. 2022), MetaCoAG (V. Mallawaarachchi and Lin 2022a, [b] 2022), and RepBin (Xue et al. 2022), where the assembly graphs are utilised in conjunction with taxonomy-independent metagenomic binning methods to recover high-quality metagenome-assembled genomes (hereafter *MAGs*) of bacterial genomes. Moreover, assembly graphs have been used for bacterial strain resolution in metagenomic data (Quince et al. 2021). However, limited studies have been conducted to resolve phage genomes in metagenomic data, particularly viral enriched metagenomes.

Computational tools have enabled large-scale studies to recover novel phages entirely from metagenomic sequencing data (Simmonds et al. 2017) and gain insights into interactions with their hosts (Nayfach, Páez-Espino, et al. 2021; M. J. Roach, McNair, et al. 2022; Hesse et al. 2022). While exciting progress has been made towards identifying new phages, viral dark matter remains vast. Current methods are either too slow or result in inaccurate or incomplete phage genomes. Generating high-quality phage genomes via *de novo* metagenome assembly is challenging due to the modular and mosaic nature of phage genomes (Lima-Mendez, Toussaint, and Leplae 2011; Hatfull 2008; Belcaid, Bergeron, and Poisson 2010). Repeat regions can result in fragmented assemblies and chimeric contigs (Casjens and Gilcrease 2009; Merrill et al. 2016). Hence, current state-of-the-art computational tools rely on the combination of either more conservative tools based on sequence- and profile-based screening (e.g. MetaPhinder (Jurtz et al. 2016)) or machine learning approaches based on nucleotide signatures (e.g. Seeker (Auslander et al. 2020), refer to Table S1 in section 1 of the Supplementary material). Resulting predictions are then evaluated using tools such as CheckV (Nayfach, Camargo, et al. 2021) and VIBRANT (Kieft, Zhou, and Anantharaman 2020) to categorise the predicted phages based on their completeness, contamination levels, and possible lifestyle (virulent or temperate) (McNair, Bailey, and Edwards 2012). Due to the supervised nature of the underlying approaches, most of these tools cannot characterise novel viruses that are significantly different from known viruses. Moreover, the approach used by these tools can be problematic with fragmented assemblies where contigs do not always represent complete genomes. In an attempt to address this limitation, tools such as MARVEL (Amgarten et al. 2018) and PHAMB (Johansen et al. 2022) were developed to identify viral metagenome-assembled genomes (vMAGs) of phages from metagenomic data. These programs rely on existing taxonomy-independent metagenomic binning tools such as MetaBAT2 (Kang et al. 2019) or VAMB (Nissen et al. 2021) and attempt to predict viral genome bins from this output using machine learning techniques.

Metagenomic binning tools are designed to capture nucleotide and sequence coverage-specific patterns of different taxonomic groups; therefore, sequences from viruses with low and uneven sequence coverage are often inaccurately binned. Many metagenomic binning tools filter out short sequences (e.g., shorter than 1,500 bp (Kang et al. 2019)), which further result in the loss of essential regions in phage genomes that are often present as short fragments in the assembly (Casjens and Gilcrease 2009). Moreover, most metagenomic binning tools struggle to distinguish viruses from genetically diverse populations with high strain diversity and quasispecies dynamics. These tools do not resolve the clustered sequences into contiguous genomes and the bins produced often contain a mixture of multiple strains resulting in poor-quality MAGs (Meyer et al. 2022). Existing solutions developed for viral quasispecies assembly only consider one species at a time (Baaijens, Stougie, and Schönhuth 2020; Freire et al. 2021, 2022) and cannot be applied to complex metagenomes. Despite the recent progress, it is challenging for currently available tools to recover complete high-quality phage genomes from metagenomic data, and a novel approach is required to address this issue. The use of connectivity information from assembly graphs could overcome these challenges (as shown in previous studies on bacterial metagenomes (V. Mallawaarachchi and Lin 2022a, [b] 2022; Lamurias et al. 2022)) to enable the recovery of high-quality phage genomes.

In this paper, we introduce Phables, a software tool that can resolve complete high-quality phage genomes from viral metagenome assemblies. First, Phables identifies phage-like components in the assembly graph using conserved genes. Second, using read mapping information, graph algorithms and flow decomposition techniques, Phables identifies the most probable combinations of varying phage genome segments within a component, leading to the recovery of accurate phage genome assemblies (Figure 1). We evaluated the quality of the resolved genomes using different assessment techniques and demonstrate that Phables produces complete and high-quality phage genomes.

**Figure 1:**
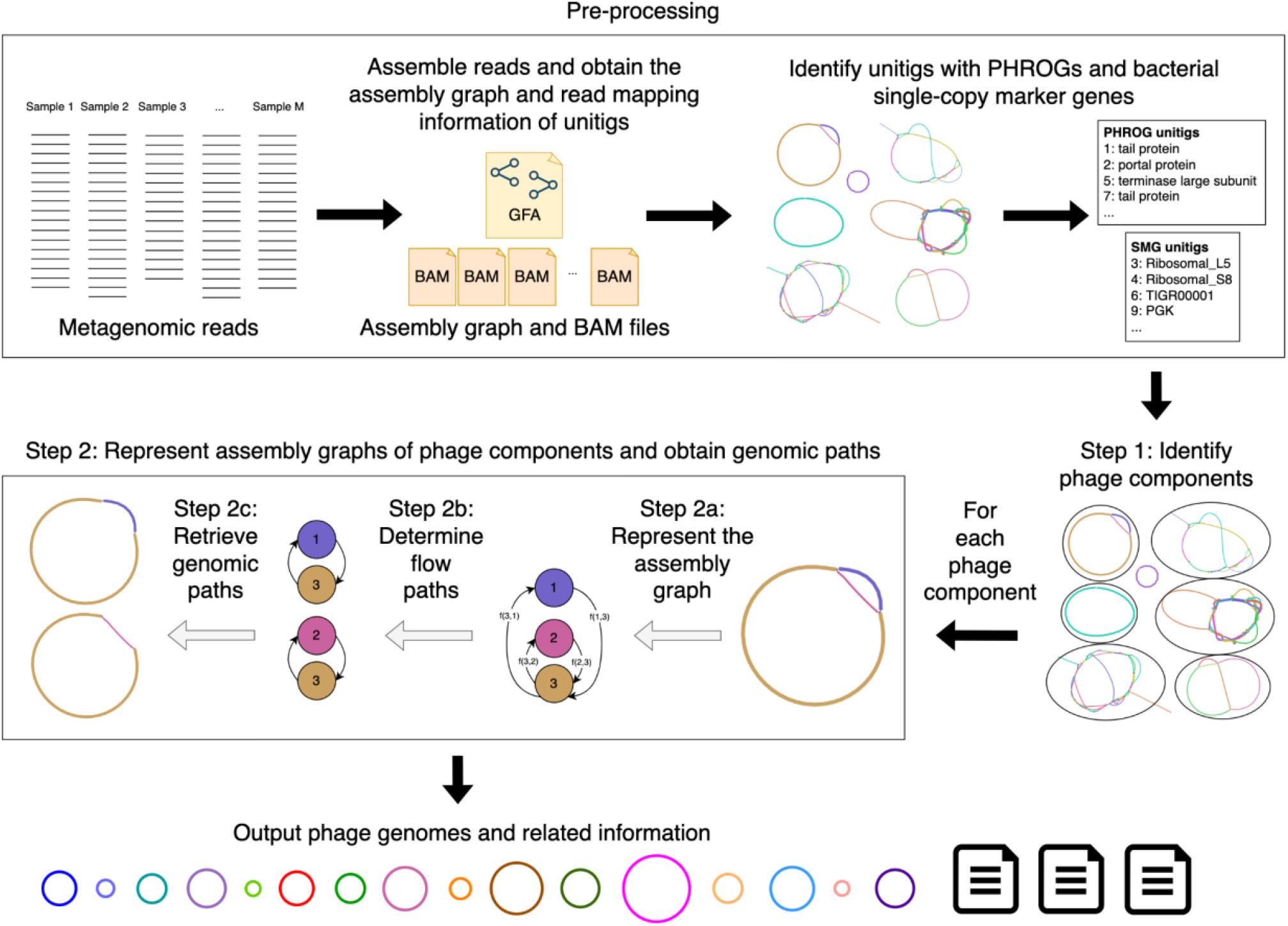
Phables workflow. Pre-processing: Assemble reads, obtain the assembly graph and read mapping information, and identify unitigs with PHROGs and bacterial single copy marker genes. Step 1: Identify phage components from the initial assembly. Step 2: For each phage component, represent the assembly graph, determine the flow paths and retrieve the genomic paths. Finally, output phage genomes and related information.

## Materials and Methods

Here we present the overall workflow of Phables (Figure 1). Metagenomic reads from single or multiple viral metagenomic samples are assembled, and the assembly graph and read mapping information are obtained. The unitig sequences from the assembly graph are extracted and screened for Prokaryotic Virus Remote Homologous Groups (PHROGs) (Terzian et al. 2021) and bacterial single-copy marker genes. Phables identifies sub-graphs (known as *phage components*) and resolves separate phage genomes from each phage component. Finally, Phables outputs the resolved phage genomes and related information. Each step of Phables is explained in detail in the following sections.

### Pre-processing

The pre-processing step performed by Phables uses an assembly graph and generates the read mapping information and the gene annotations required for Step 1 in the workflow. We recommend Hecatomb (M. J. Roach, Beecroft, et al. 2022) to assemble the reads into contigs and obtain the assembly graph. However, Phables will work with any assembly graph in Graphical Fragment Assembly (GFA) format.

The unitig sequences are extracted from the assembly graph, and the raw sequencing reads are mapped to the unitigs using Minimap2 (H. Li 2018) and Samtools (H. Li et al. 2009). Phables uses CoverM (Woodcroft and Newell 2017) to calculate the read coverage of unitigs, using the reads from all samples, and records the mean coverage (the average number of reads that map to each base of the unitig).

Phables identifies unitigs containing Prokaryotic Virus Remote Homologous Groups (PHROGs) (Terzian et al. 2021). PHROGs are viral protein families commonly used to annotate prokaryotic viral sequences. MMSeqs2 (Steinegger and Söding 2017) is used to identify PHROGs in unitigs using an identity cutoff of 30% and an e-value of less than 10^-10^ (by default).

Phables identifies unitigs containing bacterial single-copy marker genes. Most bacterial genomes have conserved genes known as single-copy marker genes (SMGs) that appear only once in a genome (Dupont et al. 2012; Albertsen et al. 2013). FragGeneScan (Rho, Tang, and Ye 2010) and HMMER (Eddy 2011) are used to identify sequences containing SMGs. SMGs are considered to be present if more than 50% (by default) of the gene length is aligned to the unitig. The list of SMGs is provided in Table S2 in section 2 of the Supplementary material.

### Step 1: Identify phage components

Phables identifies components from the final assembly graph where all of its unitigs do not have any bacterial SMGs (identified from the preprocessing step) and at least one unitig contains one or more genes belonging to a PHROG for at least one of the PHROG categories: *head and packaging, connector, tail* and *lysis* which contain known phage structural proteins and are highly conserved in tailed phages (Auslander et al. 2020) (refer to Figure S1 in section 3 of the Supplementary material for an analysis of the PHROG hits to all known phage genomes). The presence of selected PHROGs ensures the components are phage-like and represent potential phage genomes. The absence of bacterial SMGs further ensures that the components are not prophages. These identified components are referred to as *phage components*. Components that are comprised of a single circular unitig (the two ends of the unitig overlap) or a single linear unitig and that satisfy the above conditions for genes are considered *phage components* only if the unitig is longer than the predefined threshold *minlength* that is set to 2,000 bp by default, as this is the approximate lower bound of genome length for tailed phages (Luque et al. 2020).

### Step 2: Represent assembly graphs of phage components and obtain genomic paths

#### Step 2a: Represent the assembly graph

Following the definitions from STRONG (Quince et al. 2021), we define the assembly graph *G* = (*V, E*) for a phage component where *V*= {1,2,3,, | *V*| } is a collection of vertices corresponding to unitig sequences that make up a phage component and directed edges *E* ∈ *V* × *V* represent connections between unitigs. Each directed edge 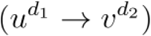 is defined by a starting vertex *u* and an ending vertex *v* (the arrow denotes the direction of the overlap), where *d*_1_,*d*_2_ ∈ {+, −}indicates whether the overlap occurs between the original sequence, indicated by a + sign or its reverse complement, indicated by a – sign.

The *weight* of each edge 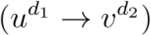 irrespective of the orientation of the edge, termed *w*_*e*_(*u* → *v*) is set to the minimum of the read coverage values of the two unitigs *u* and *v*. We also define the *confidence* of each edge 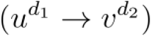 irrespective of whether the overlap occurs between the original sequence or its reverse complement, termed *c*_*e*_(*u* → *v*)is defined as the number of paired-end reads spanning across (*u* → *v*). Here, the forward read maps to unitig *u* and the reverse read maps to unitig *v*. We also define the confidence of paths (*t* →*u* → *v*) termed *c*_*p*_ (*t* →*u* → *v*) as the number of paired-end reads spanning across unitigs *t* and *u* . Paired-end information has been used in previous studies for assembling viral quasispecies (Freire et al. 2022; J. Chen, Zhao, and Sun 2018) to untangle assembly graphs. Moreover, paired-end reads are widely used in manual curation steps to join contigs from metagenome assemblies and extend them to longer sequences (L.-X. Chen et al. 2020). The more paired-end reads map to the pair of unitigs, the more confident we are about the overlap represented by the edge (refer to Figure S2 in section 4 of the Supplementary material for histograms of edge confidence).

#### Step 2b: Determine flow paths

Phables models the graph of the phage component as a flow decomposition problem and obtain the genomic paths with their coverage values calculated from the read coverages of unitigs and read mapping information. We define three cases based on the number and arrangement of unitigs present in the phage component as shown in Figure 2. Each case will be discussed in detail in the following sub-sections.

**Figure 2:**
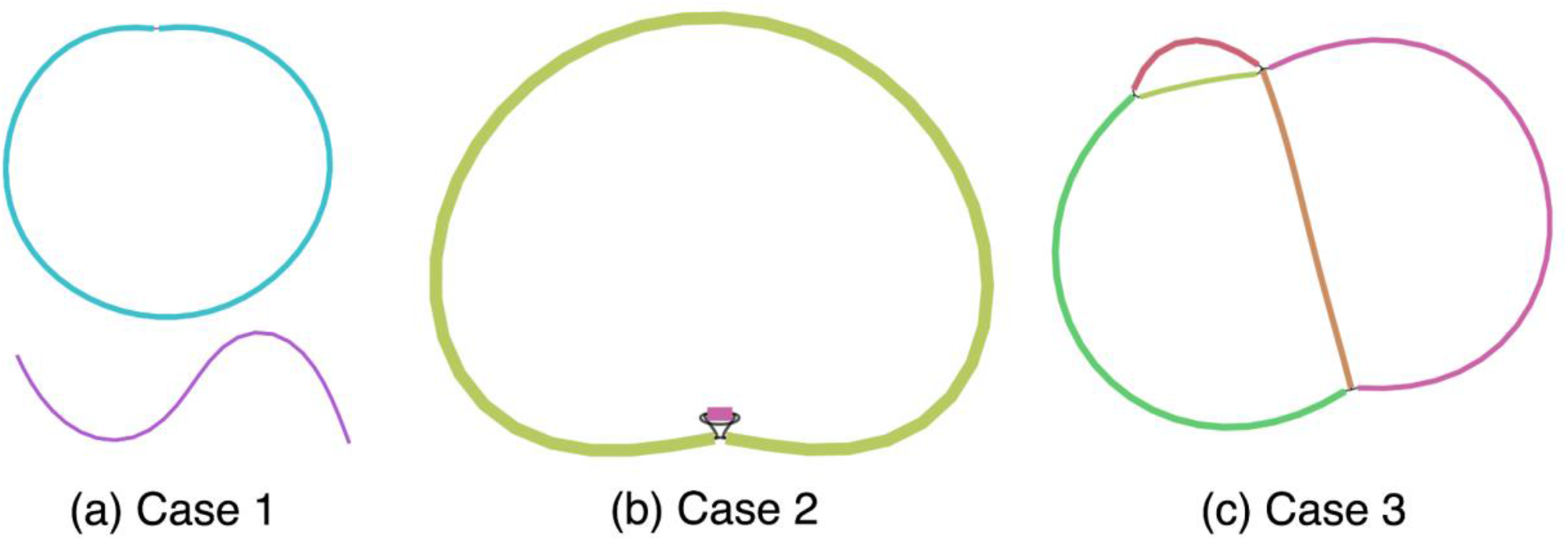
Cases of phage components. (a) Case 1 represents a phage component with one circular unitig or one linear unitig. (b) Case 2 represents a phage component with two circular unitigs connected to each other. (c) Case 3 represents a phage component that is more complex with multiple unitigs and multiple paths.

##### Case 1: Phage component consists of one circular unitig

When the phage component has only one linear/circular unitig longer than the predefined threshold *minlength*, Phables considers this unitig as one genome. The genomic path is defined as the unitig sequence itself.

##### Case 2: Phage component consists of two circular unitigs

The phage components in case 2 have two circular unitigs connected together where at least one is longer than the predefined threshold *minlength*. This is an interesting case as the shorter unitig corresponds to the terminal repeats of phages. Some phages have double-stranded repeats at their termini which are a few hundred base pairs in length and are exactly the same in every virion chromosome (i.e. they are not permuted) (Casjens and Gilcrease 2009). The terminal repeats are generated by a duplication of the repeat region in concert with packaging (Zhang and Studier 2004; Yeon-Bo Chung et al. 1990) (refer to Figure S3 in section 5 of the Supplementary material). This type of end structure could be overlooked when a phage genome sequence is determined by shotgun methods because sequence assembly can merge the two ends to give a circular sequence. Phables successfully resolves these terminal repeats to form complete genomes.

To resolve the phage component in case 2, we consider the shorter unitig (shorter than *minlength*) as the terminal repeat. Now we combine the original sequence of the terminal repeat to the beginning of the longer unitig and the reverse complement of the terminal repeat to the end of the longer unitig (refer to Figure S3 in section 5 of the Supplementary material). The coverage of the path will be set to the coverage of the longer unitig.

##### Case 3: Phage component consists of three or more unitigs

In case 3, we have more complex phage components where there are more than two unitigs forming branching paths, and we model them as a minimum flow decomposition (MFD) problem. The MFD problem decomposes a directed acyclic graph (DAG) into a minimum number of source-to-sink (*s* − *t*) paths that explain the flow values of the edges of the graph (Vatinlen et al. 2008; Dias et al. 2022). The most prominent applications of the MFD problem in bioinformatics include reconstructing RNA transcripts (Shao and Kingsford 2017; Tomescu et al. 2013; Gatter and Stadler 2019) and viral quasispecies assembly (Baaijens, Stougie, and Schönhuth 2020). The MFD problemcan be solved using integer linear programming (ILP) (Schrijver 1998).

In the viral metagenomes, we have identified structures containing several phage variant genomes, that are similar to viral quasispecies often seen in RNA viruses (Domingo and Perales 2019). Hence, Phables models each of the remaining phage components as an MFD problem and uses the MFD-ILP implementation from Dias *et al*. (Dias et al. 2022). MFD-ILP finds a *FD*(𝒫,*w*) with a set of *s* − *t* flow paths 𝒫 and associated weights *w* such that the number of flow paths is minimized. These flow paths represent possible genomic paths. An example of a phage component with possible paths is shown in Figure 3.

**Figure 3:**
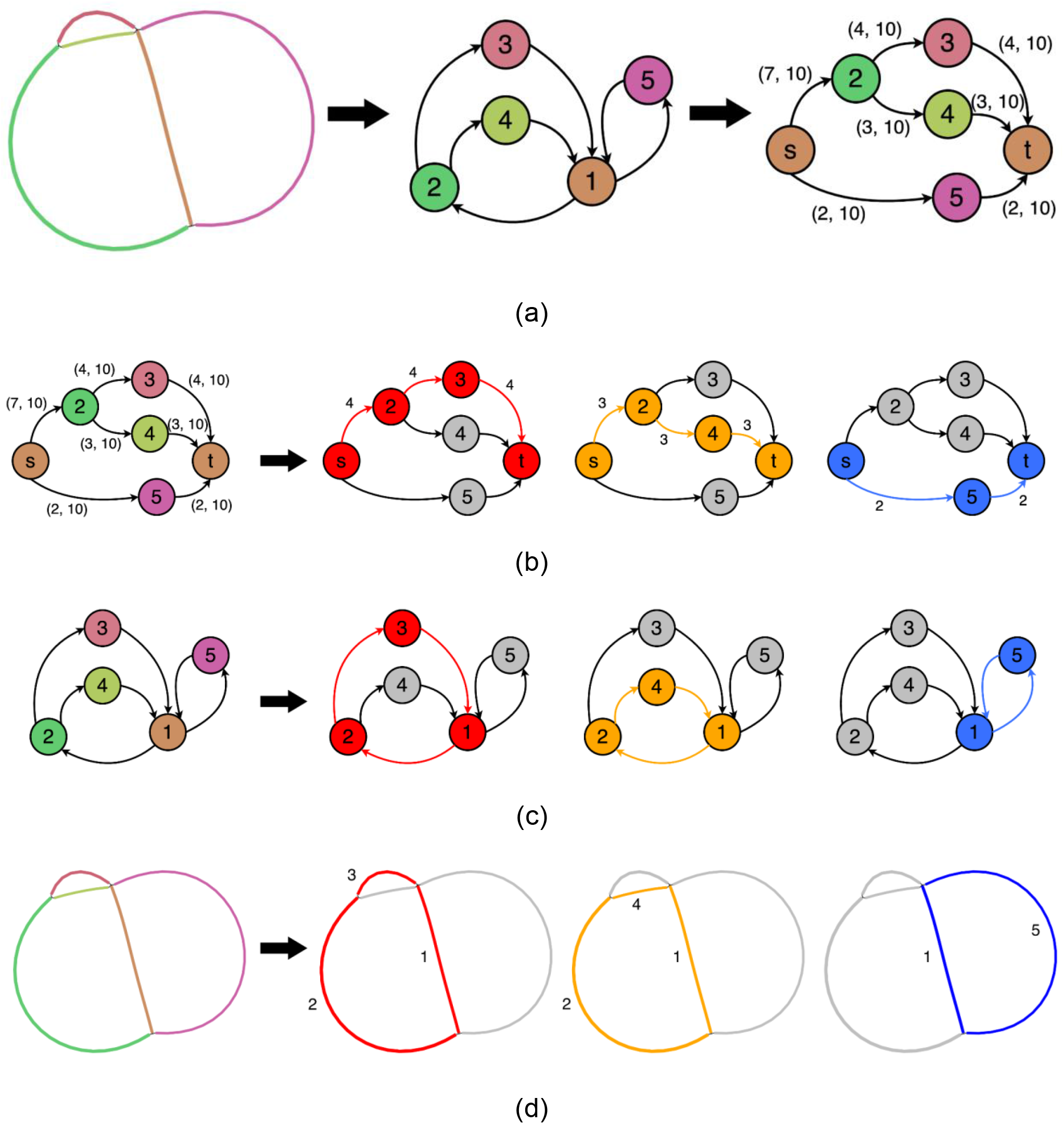
Example of a phage component (a) being modelled as a flow network and resolved into paths denoted using (b) flow network visualisation with flow values, (c) graph visualisation with directed edges and (d) Bandage (Wick et al. 2015) visualisation (with corresponding unitig numbers). Here three *s* − *t* flow paths (1→2→3, 1→2→4 and 1→5) can be obtained corresponding to three phage genomes. The arrows in (b) - (d) denote the resolution into 3 paths.

First, we convert the assembly graph of the phage component into a DAG. We start by removing *dead-ends* from *G*. We consider a vertex to be a *dead-end* if it has either no incoming edges or no outgoing edges, which arise due to errors at the start or end of reads that can create protruding chains of deviated edges (Bankevich et al. 2012). Dead-ends are particularly problematic in later steps of Phables as they can affect the continuity of genomic paths. Hence, their removal ensures that all the possible paths in the graph form closed cycles. We eliminate dead-ends by recursively removing vertices with either no incoming edges or no outgoing edges. Note that removing one dead-end can cause another vertex that is linked only to the removed one to become a dead-end, hence the removal process is done recursively.

Since a case 3 phage component forms a cyclic graph as shown in Figure 3 (a), we have to identify a vertex to represent the source/sink (referred to as *st*) in order to convert the graph to a DAG and model it as a flow network. Starting from every vertex (*source*), we conduct a breadth-first-search and identify an iterator,(*level,vertices*), where *vertices* is the non-empty list of vertices at the same distance *level* from the *source*. The method that generates this iterator is known as *bfs*_*layers* and we use the NetworkX implementation (https://networkx.org/documentation/stable/reference/algorithms/generated/networkx.algorithms.traversal.breadth_first_search.bfs_layers.html#networkx.algorithms.traversal.breadth_first_search.bfs_layers). We extract the vertices in the final layer and check if their successors are equal to *source*. If this condition holds for some vertex in *G*, we consider this vertex to be the *st* vertex of *G*. If more than one vertex satisfies the condition to be a *st* vertex, then we pick the vertex corresponding to the longest unitig as the *st* vertex. This process is carried out to find a vertex common to the flow paths (refer to Algorithm S1 in section 6 of the Supplementary material). As an example, consider vertex 1 in Figure 3 (a). When we do a breadth-first-search starting from vertex 1, the vertices in the last layer in our iterator will be 3 and 4. The successor of both 3 and 4 is vertex 1. Since the successors of the vertices in the last layer are the same as the starting vertex, we consider vertex 1 as the *st* vertex.

The edges of *G* that are weighted according to unitig coverage, may not always satisfy the conservation of flow property because of uneven sequencing depths at different regions of the genomes (Peng et al. 2012; Gunasekera et al. 2021). Hence, we use inexact flow networks which allow the edge weights to belong to an interval. Once we have identified a *st* vertex, we separate that vertex into two separate vertices for the source *s* and sink *t*. We create an inexact flow network *G*_*f*_ =(*V, E, f*, 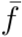) from *s* to *t* and model the rest of the vertices and edges in *G*. For example, in Figure 3 (b) vertex 1 is broken into two vertices and *s* and *t*, and the network flows from to *s* to *t*,. For every edge (*u,v*) ∈ *E* we have associated two positive integer values *f*_*uv*_ ∈ *f* and 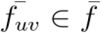, satisfying 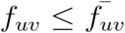 where *f*_*uv*_ = *w*_*e*_(*u* → *v*), 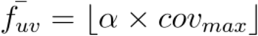, *α* ≥ 1 is the coverage multiplier parameter (1.2 by default) and *cov*_*max*_ is the maximum coverage of a unitig in the phage component. In Figure 3 (b), each edge has two values 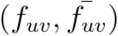 that define the flow interval for the inexact flow network *G*_*f*_. This modelling ensures that the flow through each edge is bounded by a relaxed interval between the edge weight and the maximum coverage within the component. For example, in Figure 3 (b), the (2 → 3)edge has a weight of 4 (which is the minimum of the read coverage values of the two unitigs 2 and 3 obtained from Step 2a). *α*= 1.2 and *cov*_*max*_ = 9 for the component. Hence, we set *f*_*uv*_ = 4 and 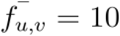.

Next, we define a set of simple paths ℛ ={*R*_1_, *R*_2_, *R*_3_,…, *R*_*l*_}where the edges that form each path have paired-end reads spanning across them, i.e. *c*_*e*_(*u* →*v*) ≥*mincov*. Enforcing these paths to contain paired-end reads ensures that genuine connections are identified and reflected in at least one decomposed path. For example, in Figure 3 (b), the edge (2 → 3) has 4 paired-end reads spanning across the edge. Hence, we add the path *R*_1_ = (2,3) to ℛ. Moreover, for a path *t* →*u* →*v* passing through the junction *u* (where the in-degree and out-degree are non-zero), we add the path *R*_*j*_ = (*t,u,v*)to ℛ, if *c*_*p*_(*t* →*u* →*v*) *≥mincov* or if |*w*_*e*_(*t* →*v*) − *w*_*e*_(*u* →*v*) | is less than a predefined threshold *cov*_*tolerance*_ (100 by default). This allows Phables to specify longer subpaths across complex junctions.

Now we model our inexact flow network *G*_*f*_ as a minimum inexact flow decomposition (MIFD) problem and determine a minimum-sized set of *s* − *t* paths 𝒫 =(*p*_1_, *p*_2_, *p*_3_,…, *p*_*k*_)and associated weights *w*= (*w*_1_,*w*_2_, *w*_3_,…,*w*_*k*_)with each *w*_*i*_ ∈ ℤ^+^ where the following conditions hold.

1. 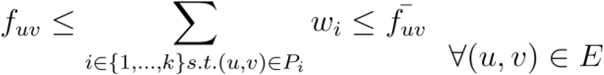

2. ∀*R*_*j*_ ∈ ℛ, *∃P*_*i*_ ∈ 𝒫such that *R*_*j*_ is a subpath of *P*_*i*_

A path *P*_*i*_ will consist of unitigs with orientation information. The weight *w*_*i*_ will be the coverage of the genome represented by the path *P*_*i*_.

#### Step 2c: Retrieve genomic paths

The flow paths obtained from cases 1 and 2 described in the previous section are directly translated to genomic paths based on the unitig sequences. In case 3, we get *s* − *t* paths from the flow decomposition step (as shown in Figure 3). The paths longer than the predefined threshold *minlength* and have a predefined coverage threshold *mincov* of (10 by default) or above are retained. For each remaining path, we remove *t* from the path as *s* and *t* are the same vertex and combine the nucleotide sequences of the unitigs corresponding to the vertices and the orientation of edges in the flow path (refer to Figure 3 (c) and (d)). Once the genomic paths of phage components are obtained, we record the constituent unitigs, path length (in bp), coverage (i.e., the flow value of the path) and the GC content of each genomic path.

### Experimental Design

#### Simulated phage dataset

We simulated reads from the following four phages with the respective read coverage values and created a simulated phage dataset (referred to as **simPhage**) to evaluate Phables.

1. Enterobacteria phage P22 (AB426868) -100×
2. Enterobacteria phage T7 (NC_001604) -150×
3. Staphylococcus phage SAP13 TA-2022 (ON911718) -200×
4. Staphylococcus phage SAP2 TA-2022 (ON911715) -400×

The Staphylococcus phage genomes have an average nucleotide identity (ANI) of 96.89%. Paired-end reads were simulated using InSilicoSeq (Gourlé et al. 2019) with the predefined MiSeq error model. We used metaSPAdes (Nurk et al. 2017) from SPAdes version 3.15.5 to assemble the reads into contigs and obtain the assembly graph for the simPhage dataset. Tables S3 and S5 in section 7 of the Supplementary material summarise the details of the simulations and assemblies.

#### Real datasets

We tested Phables on the following real viral metagenomic datasets available from the National Center for Biotechnology Information (NCBI).

1. Water samples from Nansi Lake and Dongping Lake in Shandong Province, China (NCBI BioProject number PRJNA756429), referred to as **Lake Water**
2. Soil samples from flooded paddy fields from Hunan Province, China (NCBI BioProject number PRJNA866269), referred to as **Paddy Soil**
3. Wastewater virome (NCBI BioProject number PRJNA434744), referred to as **Wastewater**
4. Stool samples from patients with IBD and their healthy household controls (NCBI BioProject number PRJEB7772) (Norman et al. 2015), referred to as **IBD**

All the real datasets were processed using Hecatomb version 1.0.1 to obtain a single assembly graph for each dataset (M. J. Roach, Beecroft, et al. 2022). Tables S3 - S5 in section 7 of the Supplementary material summarise the information about the datasets and their assemblies.

#### Tools benchmarked

We benchmarked Phables with PHAMB (Johansen et al. 2022), a viral identification tool that predicts whether MAGs represent phages and outputs genome sequences. PHAMB takes binning results from a metagenomic binning tool and predicts bins that contain bacteriophage sequences. The MAGs for PHAMB were obtained by running VAMB (version 3.0.8), a binning tool that does not rely on bacterial marker genes, in co-assembly mode on the original contigs with the author-recommended parameter `--minfasta 2000` and the `--cuda` flag. The commands used to run all the tools can be found in section 8 of the Supplementary material.

### Evaluation criteria

#### Evaluation criteria for binning tools

The resolved genomes from Phables and identified MAGs from PHAMB were evaluated using CheckV version 1.0.1 (Nayfach, Camargo, et al. 2021) (with reference database version 1.5) which compares bins against a large database of complete viral genomes. We compare the following metrics from the CheckV results.

1. CheckV viral quality
2. Completeness of sequences - number of sequences with >90% completeness
3. Contamination of sequences - number of sequences with <10% contamination
4. The number and length distribution of sequences with the following warnings
  a. Contig >1.5x longer than expected genome length
  b. High kmer_freq may indicate a large duplication

Since PHAMB predicts all viral bins, we only consider the bins from PHAMB that contain the contigs corresponding to the unitigs recovered by Phables for a fair comparison.

#### Evaluation criteria for resolved genomes

The number of components resolved by Phables for each case was recorded. The viral quality of the resolved genomes and the unitigs and contigs contained in the corresponding genomic paths were evaluated using CheckV version 1.0.1 (Nayfach, Camargo, et al. 2021). Since the reference genomes for the simPhage dataset were available, we evaluated the resolved genomes using metaQUAST (Mikheenko, Saveliev, and Gurevich 2016).

## Results

### Benchmarking results on the simulated phage dataset

We first benchmarked Phables using the simPhage dataset. We evaluated the resolved phage genomes using metaQUAST (Mikheenko, Saveliev, and Gurevich 2016). We analysed the genome coverage from metaQUAST and the coverage values reported by Phables. Figure 4 denotes the assembly graph (Bandage visualisation) of the simPhage dataset and how Phables resolved the complex case 3 component containing Staphylococcus phages.

**Figure 4:**
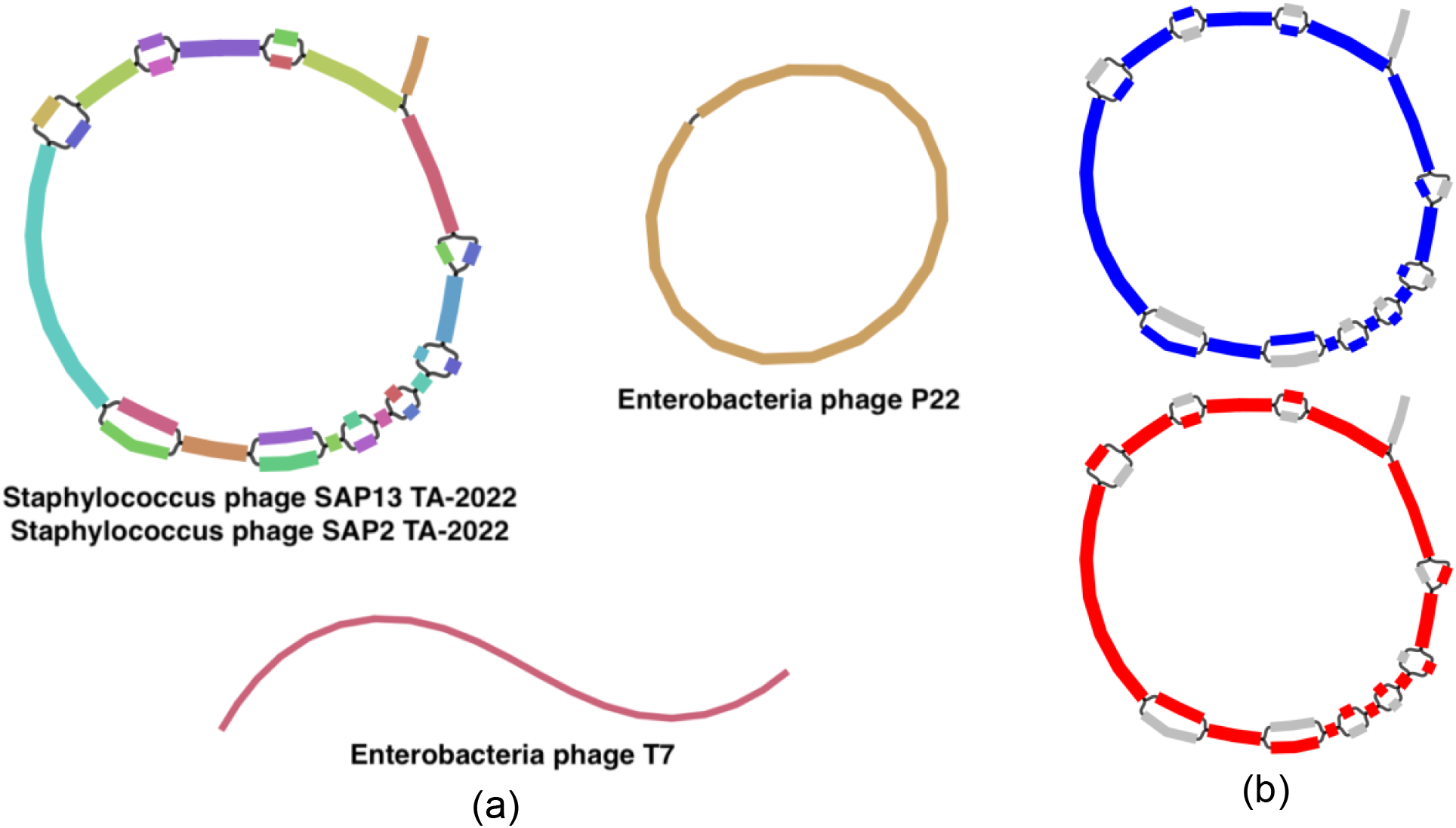
Visualisation of the (a) assembly graph from the simPhage with phage components and (b) resolution of two paths (red and blue) from the Staphylococcus phage component.

Phables recovered the two Staphylococcus phage genomes with over 92% genome completeness (refer to Table 1). The slightly low genome coverage for Staphylococcus phage SAP2 TA-2022 may have been due to the omission of the dead-end which was not properly assembled. Moreover, Phables has recovered the circular genome of Enterobacteria phage P22 and the linear genome of Enterobacteria phage T7 as well. According to Table 1, the coverage values reported from Phables are similar or close to the actual simulated coverage values of the genomes. VAMB failed to run on this dataset as there were fewer contigs than the minimum possible batch size and hence PHAMB could not be run.

**Table 1:** Evaluation results for the genomes resolved from Phables for the simPhage dataset.

| Genome | Simulated coverage | Phables predicted coverage | Genome coverage (%) |
| --- | --- | --- | --- |
| Enterobacteria phage P22 | 100 | 100 | 99.947 |
| Enterobacteria phage T7 | 150 | 150 | 99.599 |
| Staphylococcus phage SAP13 TA-2022 | 200 | 206 | 100 |
| Staphylococcus phage SAP2 TA-2022 | 400 | 401 | 92.406 |

### Benchmarking results on the real datasets

Phables resolves unitigs within phage components to produce multiple complete and high-quality genomes from the viral metagenomes (Figure 5). The genome quality of Phables results was compared with the viral-MAG prediction tool PHAMB (Johansen et al. 2022) and evaluated using CheckV (Nayfach, Camargo, et al. 2021). Figure 5 denotes the comparison of genome length distributions and genome/bin counts of different CheckV quality categories for Phables and PHAMB results. Unlike Phables, PHAMB has produced genomes with longer sequences as shown in Figures 5 (a), (c), (e) and (g), because PHAMB combines all the contigs in a bin to form one long sequence. As denoted in Figures 5 (b), (d), (f) and (h), Phables has recovered the greatest number of complete and high-quality genomes combined for all the datasets; 165 in Lake Water, 389 in Paddy Soil, 55 in Wastewater and 205 in IBD, with 49.54% more genomes recovered than PHAMB on average.

**Figure 5:**
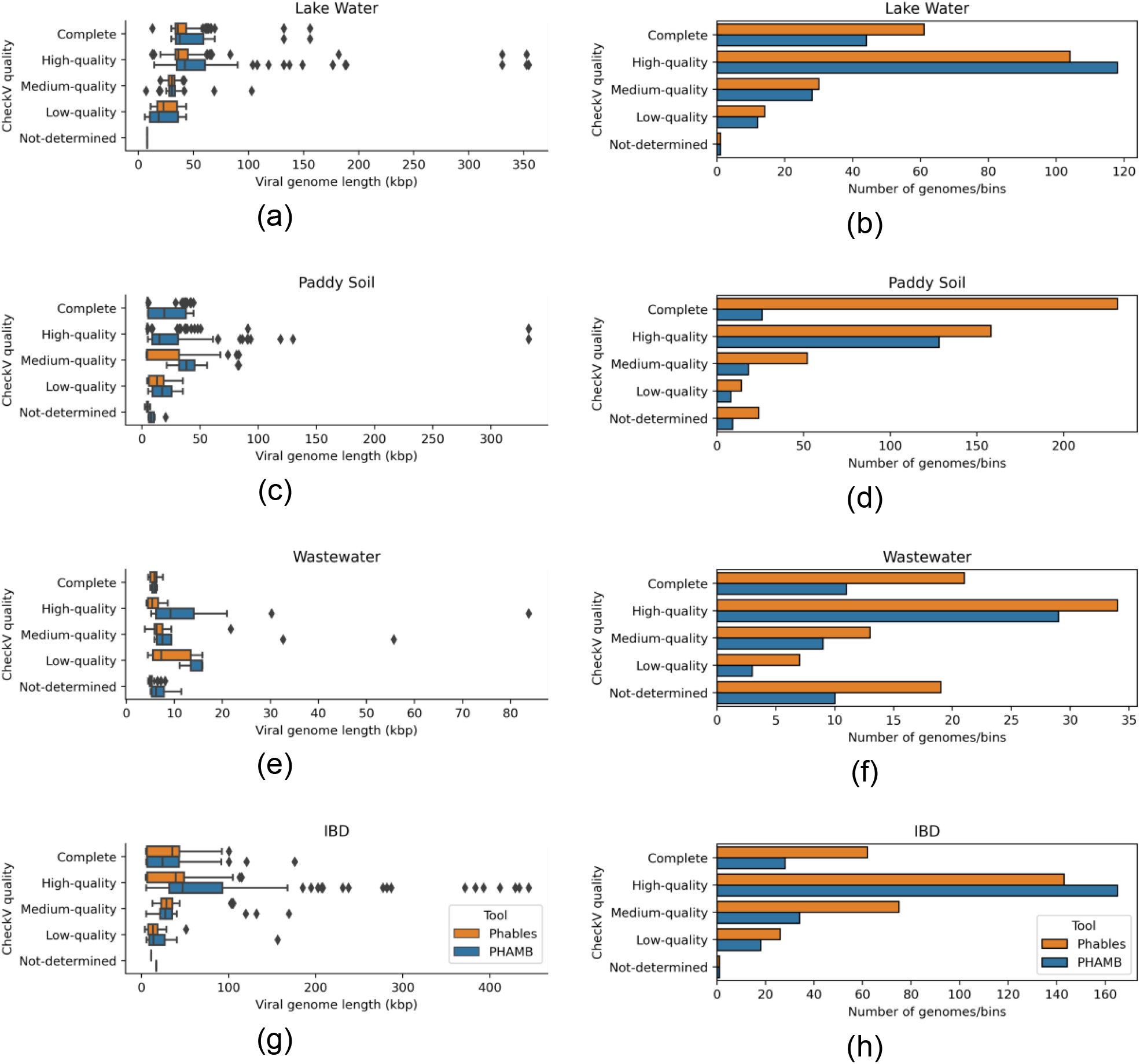
Genome length distribution (first column of figures) and abundance of genomes (second column of figures) belonging to different CheckV quality categories identified by Phables (denoted in orange) and PHAMB (Johansen et al. 2022) (denoted in blue) for the viral metagenomic datasets Lake Water, Paddy soil, Wastewater, and IBD.

Phables accurately recovers short sequences such as terminal repeats that are challenging for metagenomic binning tools to recover using the assembly graph and produces high-quality genomes. We observed that VAMB incorrectly binned the majority of the short sequences, which reduced the quality of PHAMB results. For example, the repeat sequences in the case 2 phage components identified by Phables had a mean length of 600 bp in Lake Water, 649 bp in Paddy Soil, 511 bp in Wastewater and 638 bp in IBD datasets (refer to Table S6 in section 9 of the Supplementary material for exact lengths of the sequences). All of these short sequences, except for those from the IBD dataset were found in a different bin than the bin of their connected longer sequence in the PHAMB results (8 out of 8 in Lake Water, 2 out of 2 in Paddy Soil and 1 out of 1 in Wastewater). Phables recovered these short repeat sequences along with their connected longer sequences within a phage component using the connectivity information of the assembly graph.

Phables resulted in a high number of low-quality genomes as determined by CheckV in the Wastewater dataset compared to the other datasets (Figure 5 (f)). A possible reason for this is that these may be novel phages (as they contain conserved phage markers even though CheckV categorises them as “low-quality” or “not-determined”), and so they are not yet present in the databases that CheckV relies on.

PHAMB does not carry out any resolution steps when combining the contigs of identified MAGs, which results in erroneous genome structures, high levels of contamination and duplications within genomes because of the presence of multiple variant genomes. Such duplications are identified from the warnings reported by CheckV. Hence, we evaluated the number and length distribution of sequences having CheckV warnings and the results are shown in Figure 6. PHAMB has produced the highest number of genomes with CheckV warnings and produced some very long genomes (∼355 - 485 kbp as shown in Figures 6 (a) and (g)), suggesting the combination of two or more variant genomes together in a bin. Only a few genomes produced from Phables (5 or less) contain CheckV warnings (refer to Table S8 in section 10 of the Supplementary material for the exact number of genomes with warnings). These results show that Phables accurately recovers variant genomes including regions like terminal repeats from viral metagenomic samples and produces more high-quality/complete genomes compared to existing state-of-the-art tools.

**Figure 6:**
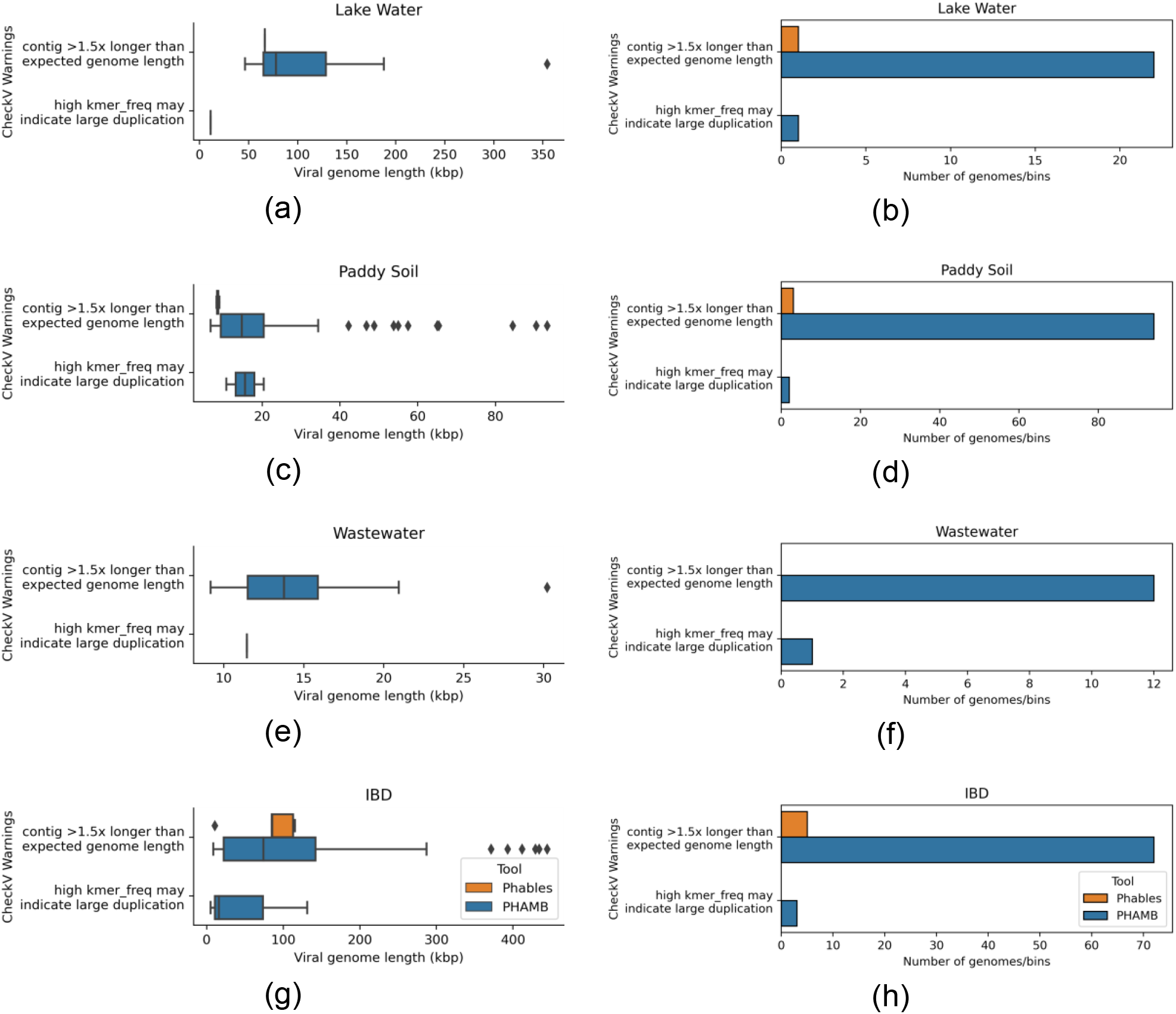
Genome length distribution (first column of figures) and abundance of genomes (second column of figures) having the selected CheckV warnings from Phables (denoted in orange) and PHAMB (Johansen et al. 2022) (denoted in blue) results for the viral metagenomic datasets Lake Water, Paddy soil, Wastewater, and IBD.

### Components resolved and comparison of resolved genomes

The number of phage components resolved by Phables under each case was recorded for all the datasets (refer to Table S7 in section 9 of the Supplementary material for the exact counts). Most of the resolved components belong to either case 1 with a single circular unitig or case 2 with the terminal repeat. When resolving case 2 components, Phables provides information regarding terminal repeats such as the length of the repeat region, that will be overlooked by other tools. Except for the IBD dataset, Phables was able to resolve all the case 3 phage components from the rest of the datasets. In a few cases, the case 3 phage components could not be resolved because Phables was unable to find a *st* vertex for these very complex bubbles (refer to Figure S8 in section 11 of the Supplementary material for examples of unresolved phage components).

Assemblers attempt to resolve longer paths in the assembly graph by connecting unitigs to form contigs (Bankevich et al. 2012; Kolmogorov et al. 2019). However, they are still unable to resolve complete genomes for complex datasets due to the mosaic nature of phage genomes and produce fragmented assemblies. Phables can be used to resolve these problematic contigs (or unitigs) and obtain high-quality genomes. Figure 7 denotes the comparison of CheckV quality of the genomes resolved in Phables and the unitigs and contigs included in the cases 2 and 3 phage components. The most complete and high-quality sequences can be found as genomes (61 and 104 for Lake Water, 231 and 158 for Paddy Soil, 21 and 34 for Wastewater, and 62 and 143 for IBD, respectively). In contrast, most medium- and low-quality genomes can be found from contigs and unitigs. Hence, genomes resolved using Phables have higher quality and will be better candidates for downstream analysis than contigs.

**Figure 7:**
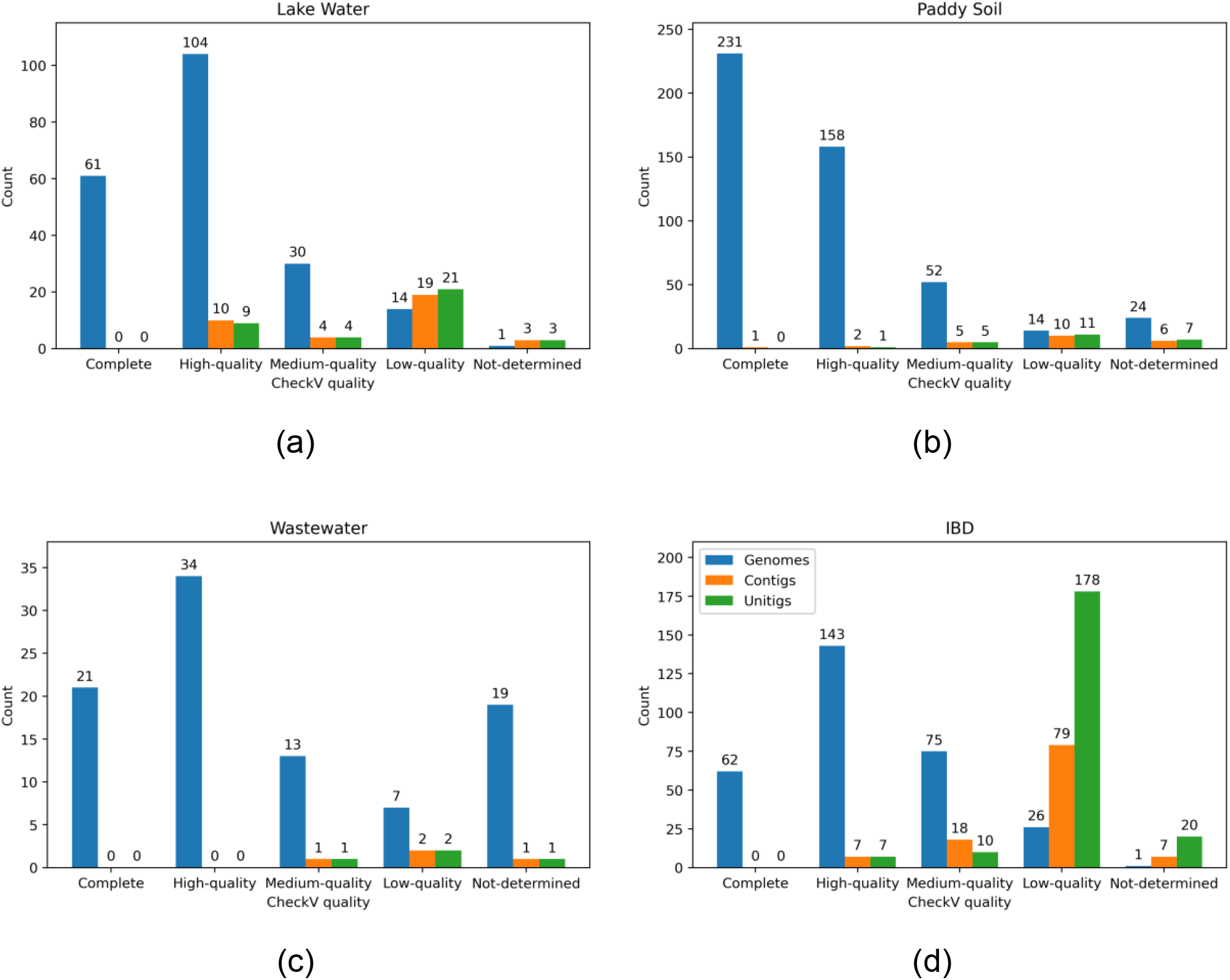
Counts of resolved genomes of Phables, unitigs and contigs included in case 2 and 3 phage components with different CheckV qualities in the viral metagenomic datasets Lake Water, Paddy soil, Wastewater, and IBD.

We compared the similarity between the genomes recovered within each case 3 phage components for the IBD dataset using *pyani* (Pritchard et al. 2015), pyGenomeViz (Shimoyama 2022) and MUMmer (Marçais et al. 2018) (refer to section 12 of the Supplementary material for the detailed results). The average nucleotide identity (ANI) analysis revealed that the genomes resolved had over 95% ANI with some genomes having over 99% ANI and over 85% alignment coverage. Moreover, as shown in Figure S10 in the Supplementary material, the mosaic genome structure can be clearly seen where some unitigs are shared between genomes and some genomes have unique unitigs. Depending on the size and location within a specific genome, these unitigs potentially correspond to functional modules. Hence, Phables can resolve highly similar variant genomes with mosaic genome structures that the assemblers are unable to distinguish.

### Phage components from other assembly methods

We extended our testing of Phables with co-assemblies obtained from other metagenome assemblers including metaSPAdes (Nurk et al. 2017) and MEGAHIT (D. Li et al. 2015) to show that the components with bubbles observed in the assembly graph are not an artefact of the assembly approach used in Hecatomb. Co-assembly is conducted by combining reads from multiple metagenomes and assembling them together, which increases the sequencing depth and provides sufficient coverage for low-abundance genomes to be recovered (Delgado and Andersson 2022). However, this becomes a computationally intensive approach as the number of samples increases, and hence we have limited the results to just the Lake Water dataset. The results are provided in section 13 of the Supplementary material and show that the phage component structures are still present in the assemblies and were correctly resolved by Phables, producing more high-quality genomes than PHAMB.

### Implementation and resource usage

The source code of Phables was implemented using Python 3.10.12 and is available as a pipeline (including all the preprocessing steps) developed using Snaketool (M. Roach et al. 2022). The commands used to run all the software can be found in section 8 of the Supplementary material. The running times of Phables core methods and including the preprocessing steps were recorded for all the datasets and can be found in Tables S10 and S11 in section 14 of the Supplementary material. The core methods of Phables can be run in under 2 minutes with less than 4 gigabytes of memory for all the datasets.

Phables uses a modified version of the MFD-ILP implementation from Dias *et al*. (Dias et al. 2022) which supports inexact flow decomposition with subpath constraints. Gurobi version 10.0.2 was used as the ILP solver. To reduce the complexity of the ILP solver, the maximum number of unitigs in a phage component to be solved was limited to 200.

## Discussion

The majority of the existing viral identification tools rely on sequence similarity- and profile-based approaches, only identifying whether assembled sequences are of viral origin, and cannot produce complete and high-quality phage genomes. Viral binning tools have been able to overcome these shortcomings up to a certain extent by producing viral MAGs, but these MAGs are fragmented and do not represent continuous genomes. Generally, the assembly process produces many short contigs where some represent regions which while important are challenging to resolve in phages, such as terminal repeat regions. These short contigs are discarded or binned incorrectly by viral binning tools, producing incomplete MAGs. Moreover, the mosaic genome structures of phage populations are a widely-documented phenomenon (Lima-Mendez, Toussaint, and Leplae 2011; Hatfull 2008; Belcaid, Bergeron, and Poisson 2010), and cannot be resolved by existing assemblers and binning tools. The resulting MAGs may contain multiple variant genomes assembled together and hence have high contamination.

Here, we introduce Phables, a new tool to resolve complete and high-quality phage genomes from viral metagenome assemblies using assembly graphs and flow decomposition techniques. We studied the assembly graphs constructed from different assembly approaches and different assembly software and consistently observed phage-like components with variation (*phage components*). Phables models the assembly graphs of these components as a minimum flow decomposition problem using read coverage and paired-end mapping information and recovers the genomic paths of different variant genomes. Experimental results confirmed that Phables recovers complete and high-quality phage genomes with mosaic genome structures, including important regions such as terminal repeats. However, Phables can identify certain plasmids as phages (e.g. *phage-plasmids* (Ravin, Svarchevsky, and Dehò 1999; Pfeifer et al. 2021; Pfeifer, Bonnin, and Rocha 2022)) because they can encode proteins homologous to phage sequences (refer to section 15 in the Supplementary material). Hence, if users run mixed-microbial communities through Phables, further downstream analysis is required to ensure that the predicted genomes do not include plasmids.

Decomposing assembly graphs has become a popular method to untangle genomes and recover variant genomes from assemblies and while we have successfully used it to obtain mostly circular phage genomes, further work needs to be conducted to handle viral metagenomes and recover the range of phage genomes. In the future, we intend to add support for long-read assemblies from dedicated metagenome assemblers that will enable Phables to enforce longer subpaths that will span across more sequences during the flow decomposition modelling. We also intend to extend the capabilities of Phables to recover linear phage genomes and explore the avenues for recovering high-quality eukaryotic viral genomes from metagenomes.

## Data and Code Availability

All the real datasets containing raw sequencing data used for this work are publicly available from their respective studies. The Lake Water dataset was downloaded from NCBI with BioProject number PRJNA756429, the Paddy Soil dataset from BioProject number PRJNA756429, the Wastewater dataset from BioProject number PRJNA434744, and the whole genome sequencing runs of the IBD data from BioProject number PRJEB7772. The sequencing reads for the simPhage dataset, all the assembled data and results from all the tools are available on Zenodo at https://zenodo.org/record/8137197.

The code of Phables is freely available on GitHub under the MIT license and can be found at https://github.com/Vini2/Phables. All analyses in this study were performed using Phables v.1.1.0 with default parameters. Phables is also available as a package on bioconda at https://anaconda.org/bioconda/phables and on PyPI at https://pypi.org/project/phables/.

## Author Contributions

V.M. designed the methods, developed the software, performed all analyses and wrote the paper. M.J.R. preprocessed the datasets. M.J.R. and P.D. assisted with developing the pipeline, optimising the steps and writing the paper. M.J.R., B.P., S.R.G., P.D., G.B. and L.K.I. tested the software and assisted with the data analysis. S.K.G., R.D.H. and A.L.K.H. curated data. All the authors reviewed the manuscript and provided detailed feedback. P.D., E.A.D. and R.A.E. conceived the project and wrote the paper with input from all authors.

## Acknowledgements

We thank Prof Christopher Quince for his insights and suggestions regarding the development of methods. This research was undertaken with the resources and services from Flinders University’s High-Performance Computing platform (“Welcome to the DeepThought HPC — DeepThought HPC Documentation” 2022).

