## Supplementary Material for "Phables: from fragmented assemblies to high-quality bacteriophage genomes"

Vijini Mallawaarachchi[1], Michael J. Roach[1], Przemyslaw Decewicz[2,1], Bhavya Papudeshi[1], Sarah K. Giles[1], Susanna R. Grigson[1], George Bouras[3], Ryan D. Hesse[1], Laura K. Inglis[1], Abbey L. K. Hutton[1], Elizabeth A. Dinsdale[1] and Robert A. Edwards[1]

[1]Flinders Accelerator for Microbiome Exploration, College of Science and Engineering, Flinders University, Bedford Park, Adelaide, SA, 5042, Australia

[2]Department of Environmental Microbiology and Biotechnology, Institute of Microbiology, Faculty of Biology, University of Warsaw, Warsaw 02-096, Poland

[3]Adelaide Medical School, The University of Adelaide, North Tce, Adelaide, SA, 5000, Australia

Keywords: bacteriophages, microbiome, metagenomics, genomics, assembly graphs

### 1. Comparison of existing tools to identify phage sequences

Table S1. Feature comparison of available tools to identify/bin sequences from phage origin in metagenomic datasets.

| Tool | Does NOT rely on a reference genome database | Uses read coverage | Uses paired-end read mapping | Uses assembly graph | Recovers short repeat regions | Does NOT need an initial binning result | Resolves high-quality and continuous genomes | Code availability | Last updated |
|---|---|---|---|---|---|---|---|---|---|
| Phables | ✓ | ✓ | ✓ | ✓ | ✓ | ✓ | ✓ | GitHub Bioconda PyPI | 2023 |
| FVE-novel (Tithi et al. 2023) | X | ✓ | ✓ | X | N/A | ✓ | ✓ | GitHub | 2020 |
| PHAMB (Johansen et al. 2022) | ✓ | ✓ | X | X | ✓* | X | X[†] | GitHub Bioconda | 2022 |
| Seeker (Auslander et al. 2020) | ✓ | X | X | X | X | N/A | X | GitHub PyPI | 2021 |
| PPR-Meta (Fang et al. 2019) | ✓ | X | X | X | ✓ | N/A | X | GitHub | 2021 |
| MARVEL (Amgarten et al. 2018) | ✓ | ✓ | X | X | X | X | X | GitHub | 2022 |
| MetaPhinder (Jurtz et al. 2016) | X | X | X | X | X | N/A | X | GitHub | 2021 |

[†] Concatenates multiple sequences without being resolved in the proper order to form a single genome.

* May not correctly associate short repeats with the relevant genomes/MAGs.

N/A: not applicable or not mentioned

#### 2. Bacterial single-copy marker genes

Table S2. List of bacterial single-copy marker genes used to filter components.

| Name | Accession | Description |
|---|---|---|
| PGK | PF00162.14 | Phosphoglycerate kinase |
| Ribosomal_L23 | PF00276.15 | Ribosomal protein L23 |
| Ribosomal_L5 | PF00281.14 | Ribosomal protein L5 |
| Ribosomal_L3 | PF00297.17 | Ribosomal protein L3 |
| Ribosomal_L6 | PF00347.18 | Ribosomal protein L6 |
| Ribosomal_S17 | PF00366.15 | Ribosomal protein S17 |
| Ribosomal_S9 | Ribosomal_S9 | Ribosomal protein S9/S16 |
| Ribosomal_S8 | PF00410.14 | Ribosomal protein S8 |
| Ribosomal_S11 | PF00411.14 | Ribosomal protein S11 |
| Ribosomal_S13 | PF00416.17 | Ribosomal protein S13/S18 |
| Ribosomal_L10 | PF00466.15 | Ribosomal protein L10 |
| Ribosomal_L4 | PF00573.17 | Ribosomal protein L4/L1 family |
| tRNA-synt_1d | PF00750.14 | tRNA synthetases class I (R) |
| GrpE | PF01025.14 | GrpE |
| Methyltransf_5 | PF01795.14 | MraW methylase family |
| TIGR00001 | TIGR00001 | rpmI_bact: ribosomal protein L35 |
| TIGR00002 | TIGR00002 | S16: ribosomal protein S16 |
| TIGR00009 | TIGR00009 | L28: ribosomal protein L28 |
| TIGR00012 | TIGR00012 | L29: ribosomal protein L29 |
| TIGR00019 | TIGR00019 | prfA: peptide chain release factor 1 |
| TIGR00029 | TIGR00029 | S20: ribosomal protein S20 |
| TIGR00043 | TIGR00043 | TIGR00043: probable rRNA maturation factor YbeY |

| | | |
|---|---|---|
| TIGR00059 | TIGR00059 | L17: ribosomal protein L17 |
| TIGR00060 | TIGR00060 | L18_bact: ribosomal protein L18 |
| TIGR00061 | TIGR00061 | L21: ribosomal protein L21 |
| TIGR00062 | TIGR00062 | L27: ribosomal protein L27 |
| TIGR00064 | TIGR00064 | ftsY: signal recognition particle-docking protein FtsY |
| TIGR00082 | TIGR00082 | rbfA: ribosome-binding factor A |
| TIGR00086 | TIGR00086 | smpB: SsrA-binding protein |
| TIGR00092 | TIGR00092 | TIGR00092: GTP-binding protein YchF |
| TIGR00115 | TIGR00115 | tig: trigger factor |
| TIGR00116 | TIGR00116 | tsf: translation elongation factor Ts |
| TIGR00152 | TIGR00152 | TIGR00152: dephospho-CoA kinase |
| TIGR00158 | TIGR00158 | L9: ribosomal protein L9 |
| TIGR00165 | TIGR00165 | S18: ribosomal protein S18 |
| TIGR00166 | TIGR00166 | S6: ribosomal protein S6 |
| TIGR00168 | TIGR00168 | infC: translation initiation factor IF-3 |
| TIGR00234 | TIGR00234 | tyrS: tyrosine--tRNA ligase |
| TIGR00337 | TIGR00337 | PyrG: CTP synthase |
| TIGR00344 | TIGR00344 | alaS: alanine--tRNA ligase |
| TIGR00362 | TIGR00362 | DnaA: chromosomal replication initiator protein DnaA |
| TIGR00388 | TIGR00388 | glyQ: glycine--tRNA ligase, alpha subunit |
| TIGR00389 | TIGR00389 | glyS_dimeric: glycine--tRNA ligase |
| TIGR00392 | TIGR00392 | ileS: isoleucine--tRNA ligase |
| TIGR00396 | TIGR00396 | leuS_bact: leucine--tRNA ligase |
| TIGR00408 | TIGR00408 | proS_fam_I: proline--tRNA ligase |
| TIGR00409 | TIGR00409 | proS_fam_II: proline--tRNA ligase |
| TIGR00414 | TIGR00414 | serS: serine--tRNA ligase |
| TIGR00418 | TIGR00418 | thrS: threonine--tRNA ligase |

| | | |
|---|---|---|
| TIGR00420 | TIGR00420 | trmU: tRNA (5-methylaminomethyl-2-thiouridylate)-methyltransferase |
| TIGR00422 | TIGR00422 | valS: valine--tRNA ligase |
| TIGR00435 | TIGR00435 | cysS: cysteine--tRNA ligase |
| TIGR00436 | TIGR00436 | era: GTP-binding protein Era |
| TIGR00442 | TIGR00442 | hisS: histidine--tRNA ligase |
| TIGR00459 | TIGR00459 | aspS_bact: aspartate--tRNA ligase |
| TIGR00460 | TIGR00460 | fmt: methionyl-tRNA formyltransferase |
| TIGR00468 | TIGR00468 | pheS: phenylalanine--tRNA ligase, alpha subunit |
| TIGR00471 | TIGR00471 | pheT_arch: phenylalanine--tRNA ligase, beta subunit |
| TIGR00472 | TIGR00472 | pheT_bact: phenylalanine--tRNA ligase, beta subunit |
| TIGR00487 | TIGR00487 | IF-2: translation initiation factor IF-2 |
| TIGR00496 | TIGR00496 | frr: ribosome recycling factor |
| TIGR00575 | TIGR00575 | dnlj: DNA ligase, NAD-dependent |
| TIGR00631 | TIGR00631 | uvrb: excinuclease ABC subunit B |
| TIGR00663 | TIGR00663 | dnan: DNA polymerase III, beta subunit |
| TIGR00755 | TIGR00755 | ksgA: dimethyladenosine transferase |
| TIGR00810 | TIGR00810 | secG: preprotein translocase, SecG subunit |
| TIGR00855 | TIGR00855 | L12: ribosomal protein L7/L12 |
| TIGR00922 | TIGR00922 | nusG: transcription termination/antitermination factor NusG |
| TIGR00952 | TIGR00952 | S15_bact: ribosomal protein S15 |
| TIGR00959 | TIGR00959 | ffh: signal recognition particle protein |
| TIGR00963 | TIGR00963 | secA: preprotein translocase, SecA subunit |
| TIGR00964 | TIGR00964 | secE_bact: preprotein translocase, SecE subunit |
| TIGR00967 | TIGR00967 | 3a0501s007: preprotein translocase, SecY subunit |
| TIGR00981 | TIGR00981 | rpsL_bact: ribosomal protein S12 |
| TIGR01009 | TIGR01009 | rpsC_bact: ribosomal protein S3 |

| | | |
|---|---|---|
| TIGR01011 | TIGR01011 | rpsB_bact: ribosomal protein S2 |
| TIGR01017 | TIGR01017 | rpsD_bact: ribosomal protein S4 |
| TIGR01021 | TIGR01021 | rpsE_bact: ribosomal protein S5 |
| TIGR01024 | TIGR01024 | rplS_bact: ribosomal protein L19 |
| TIGR01029 | TIGR01029 | rpsG_bact: ribosomal protein S7 |
| TIGR01030 | TIGR01030 | rpmH_bact: ribosomal protein L34 |
| TIGR01031 | TIGR01031 | rpmF_bact: ribosomal protein L32 |
| TIGR01032 | TIGR01032 | rplT_bact: ribosomal protein L20 |
| TIGR01044 | TIGR01044 | rplV_bact: ribosomal protein L22 |
| TIGR01049 | TIGR01049 | rpsJ_bact: ribosomal protein S10 |
| TIGR01050 | TIGR01050 | rpsS_bact: ribosomal protein S19 |
| TIGR01059 | TIGR01059 | gyrB: DNA gyrase, B subunit |
| TIGR01063 | TIGR01063 | gyrA: DNA gyrase, A subunit |
| TIGR01066 | TIGR01066 | rplM_bact: ribosomal protein L13 |
| TIGR01067 | TIGR01067 | rplN_bact: ribosomal protein L14 |
| TIGR01071 | TIGR01071 | rplO_bact: ribosomal protein L15 |
| TIGR01079 | TIGR01079 | rplX_bact: ribosomal protein L24 |
| TIGR01164 | TIGR01164 | rplP_bact: ribosomal protein L16 |
| TIGR01169 | TIGR01169 | rplA_bact: ribosomal protein L1 |
| TIGR01171 | TIGR01171 | rplB_bact: ribosomal protein L2 |
| TIGR01391 | TIGR01391 | dnaG: DNA primase |
| TIGR01393 | TIGR01393 | lepA: GTP-binding protein LepA |
| TIGR01632 | TIGR01632 | L11_bact: ribosomal protein L11 |
| TIGR01953 | TIGR01953 | NusA: transcription termination factor NusA |
| TIGR02012 | TIGR02012 | tigrfam_recA: protein RecA |
| TIGR02013 | TIGR02013 | rpoB: DNA-directed RNA polymerase, beta subunit |
| TIGR02027 | TIGR02027 | rpoA: DNA-directed RNA polymerase, alpha subunit |

| | | |
|---|---|---|
| TIGR02191 | TIGR02191 | RNaseIII: ribonuclease III |
| TIGR02350 | TIGR02350 | prok_dnaK: chaperone protein DnaK |
| TIGR02386 | TIGR02386 | rpoC_TIGR: DNA-directed RNA polymerase, beta' subunit |
| TIGR02387 | TIGR02387 | rpoC1_cyan: DNA-directed RNA polymerase, gamma subunit |
| TIGR02397 | TIGR02397 | dnaX_nterm: DNA polymerase III, subunit gamma and tau |
| TIGR02432 | TIGR02432 | lysidine_TilS_N: tRNA(Ile)-lysidine synthetase |
| TIGR02729 | TIGR02729 | Obg_CgtA: Obg family GTPase CgtA |
| TIGR03263 | TIGR03263 | guanyl_kin: guanylate kinase |
| TIGR03594 | TIGR03594 | GTPase_EngA: ribosome-associated GTPase EngA |

#### 3. Analysis of PHROG annotations in all known phage genomes

We annotated all the 25,596 complete phage genomes available from the INfrastructure for a PHAge REference Database (INPHARED) (Cook et al. 2021) (as of the May 2023 update) using Prokaryotic Virus Remote Homologous Groups (PHROGs annotation v4). We then selected the annotations belonging to the PHROG categories: *head and packaging*, *connector*, *tail* and *lysis*. Out of 109,404 PHROGs, a total of 2705 PHROGs were selected (head and packaging: 989, connector: 146, tail: 1256 and lysis: 314). Figure S1 denotes the histogram of counts of PHROGs in the phage genomes. The annotation was done using MMSeqs2 (Steinegger and Söding 2017). Figure S1 denotes the histogram of counts of PHROGs in the phage genomes.

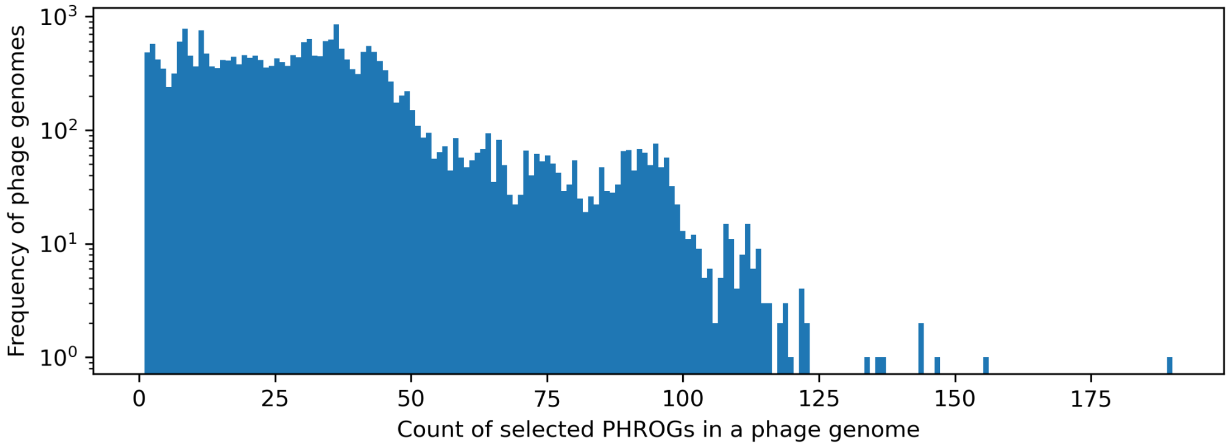


Figure S1. Histograms of counts of PHROGs found in the 25,596 complete phage genomes available from the INfrastructure for a PHAge REference Database.

Using the PHROG hits from the selected categories, we found that,

- 24,533 phage genomes (95.85%) have at least one PHROG hit.
- 24,050 phage genomes (93.96%) have two or more PHROG hits.
- 22,706 phage genomes (88.71%) have five or more PHROG hits.
- 20,319 phage genomes (79.38%) have ten or more PHROG hits.
- 15,152 phage genomes (59.20%) have at least one PHROG from all the categories.
- 24,131 phage genomes (94.28%) have at least one head and packaging PHROG.
- 16,011 phage genomes (62.55%) have at least one connector PHROG.
- 23,162 phage genomes (90.49%) have at least one tail PHROG.
- 21,081 phage genomes (82.36%) have at least one lysis PHROG.

### 4. Edge confidence

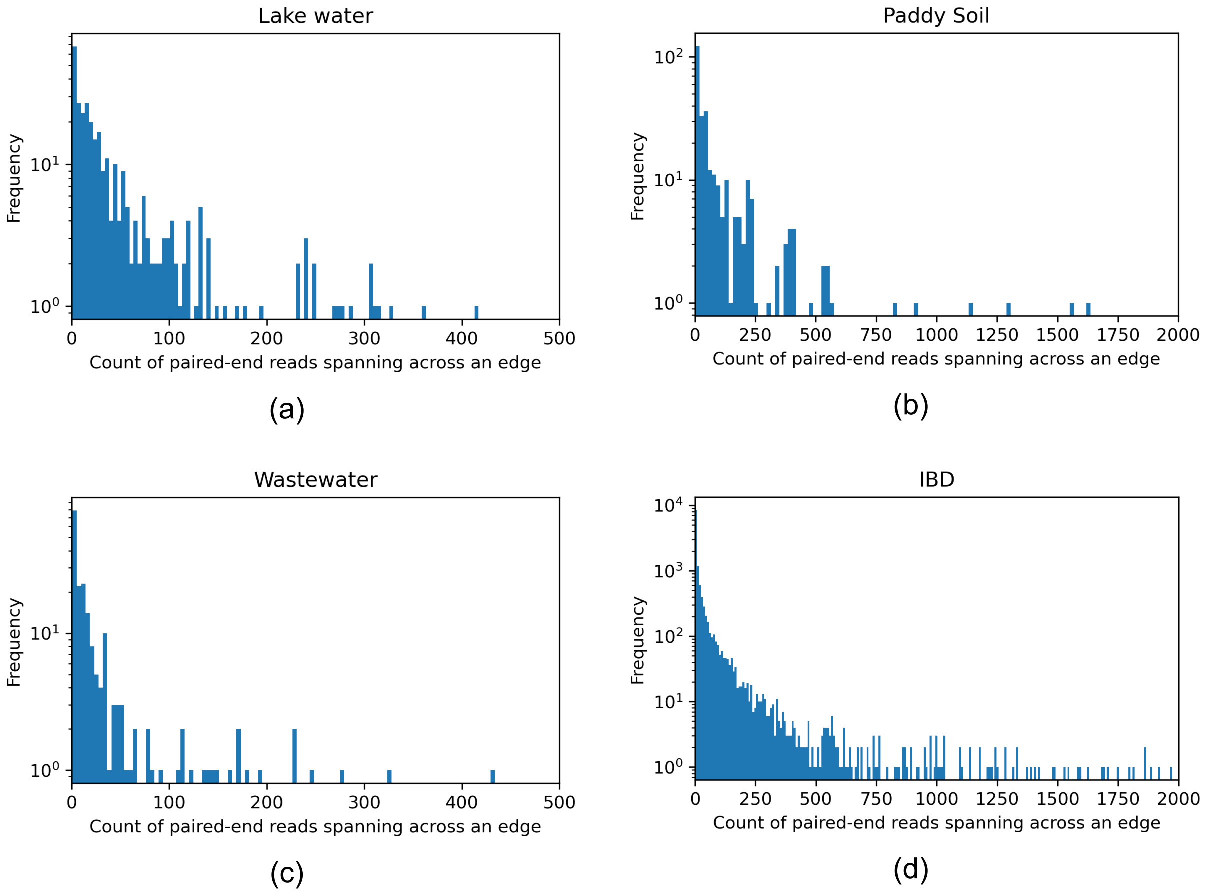


Figure S2. Histograms of paired-end read counts spanning across edges in the assembly graph (edge confidence) for the metagenomic datasets (a) Lake Water, (b) Paddy soil, (c) Wastewater, and (d) IBD.

#### 5. Resolving a case 2 phage component

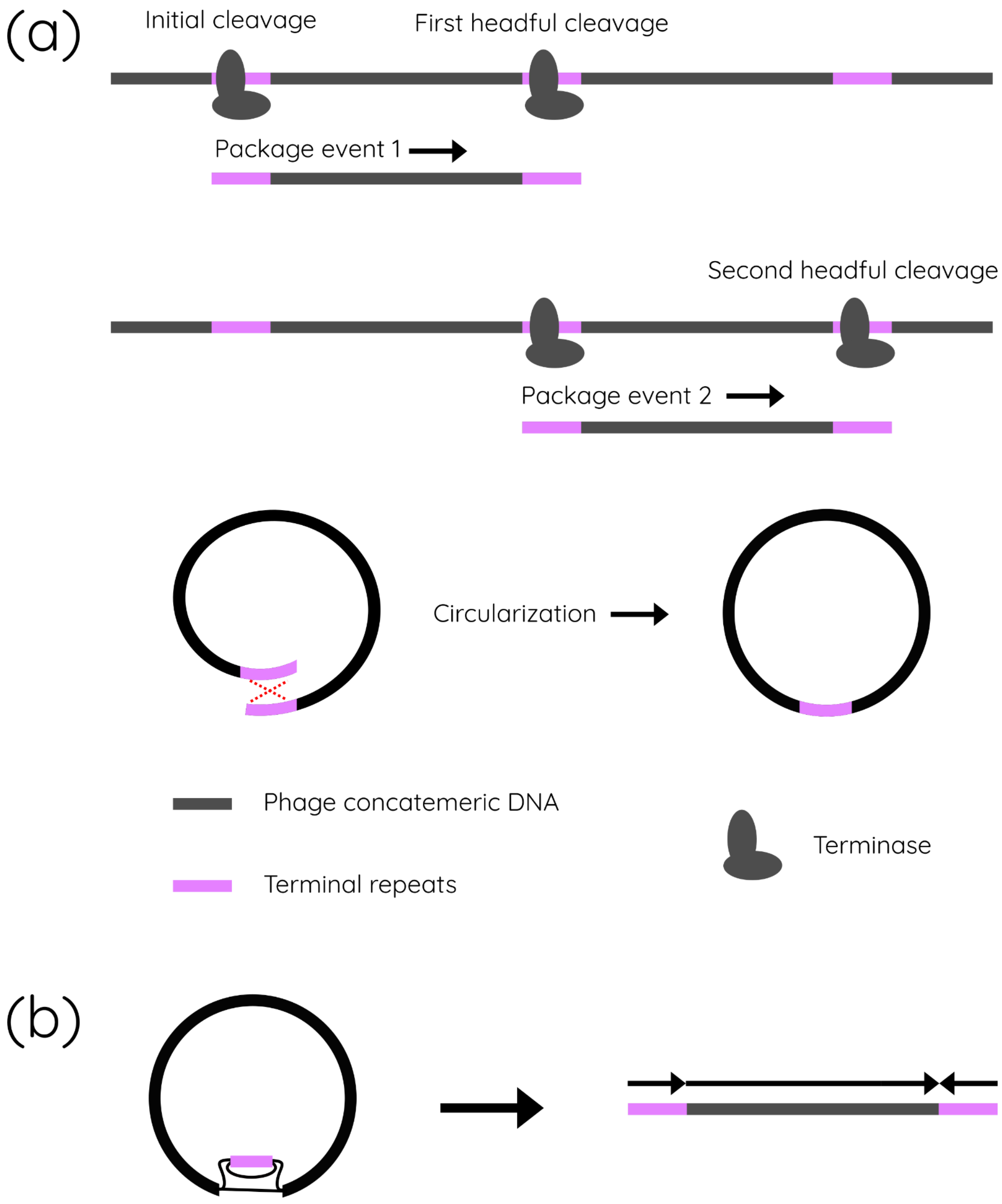


Figure S3: Example of resolving a case 2 phage component. (a) Each packaging event starts and finishes at a packaging recognition site that gets duplicated. The duplicated regions recombine and collapse during circularisation to form a continuous circular genome. (b) Representation of a case 2 phage component with a terminal repeat assembled separately and then resolved into a linearised genome.

#### 6. Algorithms

Algorithm S1: Identify $st$ vertex of $G$

Input: Assembly graph for a phage bubble ($G$)
Output: $st$, the source/sink vertex
Function: $getSourceSink(G)$

```
    st_nodes ← ∅
    forall source ∈ G.vertices do
        source_is_st ← True
        iteratore ← bfs_layers(G, source)
        last_layer ← list of vertices in the last layer of iterator
        forall v ∈ last_layer do
            v_successors ← G.successor(v)
            if v_successors ≠ ∅ and source ∉ v_successors then
                source_is_st ← False
                break
            if v_successors = ∅ then
                source_is_st ← False
        if source_is_st then
            st_nodes ← st_nodes ∪ source
    if st_nodes ≠ ∅ then
        return arg max_{length} st_node
    else
        return False
```

### 7. Information about the datasets

Table S3. Information about the simPhage dataset

| Phage genome | Accession number | Simulated coverage (x) |
|---|---|---|
| Enterobacteria phage P22 | AB426868 | 100 |
| Enterobacteria phage T7 | NC_001604 | 150 |
| Staphylococcus phage SAP13 TA-2022 | ON911718 | 200 |
| Staphylococcus phage SAP2 TA-2022 | ON911715 | 400 |

Table S4. Accession numbers of the runs used for each real dataset

| Dataset | BioProject | SRA accession numbers |
|---|---|---|
| Lake Water | PRJNA756429 | SRR15557808, SRR15557809, SRR15557810, SRR15557811, SRR15557812, SRR15557813 |
| Paddy Soil | PRJNA756429 | SRR20958178, SRR20958179, SRR20958180, SRR20958181, SRR20958182, SRR2095818 |
| Wastewater | PRJNA434744 | SRR6872420, SRR6872421, SRR6872422, SRR6872423, SRR6872424, SRR6872425, SRR6872426, SRR6872427, SRR6872428, SRR6872429, SRR6872430, SRR6872431, SRR6872432, SRR6872433, SRR6872434, SRR6872435, SRR6872436, SRR6872437 |
| IBD (Norman et al. 2015) | PRJEB7772 | ERR843912, ERR843913, ERR843914, ERR843915, ERR843916, ERR843917, ERR843918, ERR843919, ERR843920, ERR843921, ERR843922, ERR843923, ERR843924, ERR843925, ERR843926, ERR843927, ERR843928, ERR843929, ERR843930, ERR843931, ERR843932, ERR843933, ERR843934, ERR843935, ERR843936, ERR843937, ERR843938, ERR843939, ERR843940, ERR843941, ERR843942, ERR843943, ERR843944, ERR843945, ERR843946, ERR843947, ERR843948, ERR843949, ERR843950, ERR843951, ERR843952, ERR843953, ERR843954, ERR843955, ERR843956, ERR843957, ERR843958, ERR843959, ERR843960, ERR843961, ERR843962, ERR843963, ERR843964, ERR843965, ERR843966, ERR843967, ERR843968, ERR843969, ERR843970, ERR843971, ERR843972, ERR843973, ERR843974, ERR843975, ERR843976, ERR843977, ERR843978, ERR843979, ERR843980, ERR843981, ERR843982, ERR843983, |

| | | |
|---|---|---|
| | | ERR843984, ERR843985, ERR843986, ERR843987, ERR843988, ERR843989, ERR843990, ERR843991, ERR843992, ERR843993, ERR843994, ERR843995, ERR843996, ERR843997, ERR843998, ERR843999, ERR844000, ERR844001, ERR844002, ERR844003, ERR844004, ERR844005, ERR844006, ERR844007, ERR844008, ERR844009, ERR844010, ERR844011, ERR844012, ERR844013, ERR844014, ERR844015, ERR844016, ERR844017, ERR844018, ERR844019, ERR844020, ERR844021, ERR844022, ERR844023, ERR844024, ERR844025, ERR844026, ERR844027, ERR844028, ERR844029, ERR844030, ERR844031, ERR844032, ERR844033, ERR844034, ERR844035, ERR844036, ERR844037, ERR844038, ERR844039, ERR844040, ERR844041, ERR844042, ERR844043, ERR844044, ERR844045, ERR844046, ERR844047, ERR844048, ERR844049, ERR844050, ERR844051, ERR844052, ERR844053, ERR844054, ERR844055, ERR844056, ERR844057, ERR844058, ERR844059, ERR844060, ERR844061, ERR844062, ERR844063, ERR844064, ERR844065, ERR844066, ERR844067, ERR844068, ERR844069, ERR844070, ERR844071, ERR844072, ERR844073, ERR844074, ERR844075, ERR844076, ERR844077, ERR844078, ERR844079, ERR844080, ERR844081 |

Table S5. Information about the assemblies of all the datasets

| Dataset | No. of samples | Read length (bp) | Assembly size (Mb) | Number of contigs | Number of unitigs | Number of edges |
|---|---|---|---|---|---|---|
| simPhage | 1 | 2 × 300 | 161.19 | 7 | 17 | 19 |
| Lake Water | 6 | 2 × 300 | 282.52 | 129 107 | 129 212 | 702 |
| Paddy Soil | 6 | 2 × 150 | 630.03 | 248 941 | 249 193 | 2226 |
| Wastewater | 18 | 2 × 300 | 53.62 | 22 035 | 22 081 | 135 |
| IBD | 171 | 2 × 250 | 106.46 | 23 732 | 26 329 | 35 726 |
| Lake Water - metaSPAdes | 6 | 2 × 300 | 234.88 | 344 873 | 887 214 | 1 121 225 |
| Lake Water - MEGAHIT | 6 | 2 × 300 | 633.52 | 1 200 868 | N/A* | 850 422 |

* Assembly graphs of MEGAHIT assemblies contain the contig sequences.

#### 8. Commands used

##### Hecatomb

```
> hecatomb run --reads reads/ --threads 16
```

##### metaSPAdes

```
> spades.py --meta -1 reads_R1.fastq -2 reads_R2.fastq -o
metaspades_output -t 16 -m 500
```

##### MEGAHIT

```
# Run MEGAHIT
> megahit -1 reads_R1.fastq -2 reads_R2.fastq -o MEGAHIT -t 16

# Generate FASTG file
> megahit_toolkit contig2fastg 141 final.contigs.fa >
final.contigs.fastg

# Convert FASTG file to GFA
# Using https://github.com/lh3/gfa1/blob/master/misc/fastg2gfa.c
> fastg2gfa final.contigs.fastg > final.contigs.gfa
```

##### VAMB

```
# Get BAM files
> minimap2 -t 16 -N 50 -ax sr assembly.fasta
reads_sample_n_R1.fastq.gz reads_sample_n_R2.fastq.gz | samtools
view -F 3584 -b --threads 16 > bam_files/reads_sample_n.bam

# Run VAMB
> vamb --cuda --outdir vamb --fasta assembly.fasta --bamfiles
bam_files/*.bam --minfasta 2000
```

##### PHAMB

```
# Run DeepVirFinder and move results
> python3 ~/DeepVirFinder/dvf.py -i assembly.fasta -o DVF -l
2000 -c 1
> mv DVF/contigs.fna_gt2000bp_dvfpred.txt
annotations/all.DVF.predictions.txt
```

```
# Run Prodigal
> prodigal -i assembly.fasta -d genes.fna -a proteins.faa -p
meta -g 11

# Run hmmsearch
> hmmsearch --cpu {threads} -E 1.0e-05 -o output.txt --tblout
annotations/all.hmmMiComplete105.tbl <micompleteDB> proteins.faa
> hmmsearch --cpu {threads} -E 1.0e-05 -o output.txt --tblout
annotations/all.hmmVOG.tbl <VOGDB> proteins.faa

# Run PHAMB
> run_RF.py assembly.fasta vamb/clusters.tsv annotations
PHAMB_resultdir
```

##### Phables

```
> phables run --input assembly_graph.gfa --reads fastq_reads
--threads 16
```

##### CheckV

```
> checkv end_to_end all_sequences.fasta checkv_output -t 16
```

##### metaQUAST

```
> metaquast.py -R refs/ -o metaquast_output --labels
resolved_paths.fasta --threads 16
```

#### 9. Further details about resolved components

Table S6. Length of sequences in case 2 components resolved by Phables

| Dataset | Component | Length of longer sequence (bp) | Length of shorter sequence (bp) |
|---|---|---|---|
| Lake Water | 1 | 41 101 | 525 |
| | 2 | 34 378 | 618 |
| | 3 | 31 972 | 672 |
| | 4 | 31 043 | 522 |
| | 5 | 60 221 | 504 |
| | 6 | 37 094 | 944 |
| | 7 | 32 944 | 505 |
| | 8 | 33 114 | 508 |
| | Mean | 37 734 | 600 |
| Paddy Soil | 1 | 3327 | 713 |
| | 2 | 50 917 | 585 |
| | Mean | 27 122 | 649 |
| Wastewater | 1 | 7930 | 511 |
| | Mean | 7930 | 511 |
| IBD | 1 | 4310 | 566 |
| | 2 | 3566 | 709 |
| | Mean | 3938 | 638 |

Table S7. The number of components found and resolved for different cases of phage components

| Dataset | Case 1 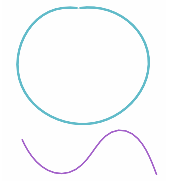 | Case 2 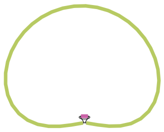 | Cae 3 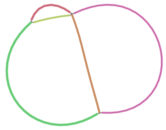 | |
|---|---|---|---|---|
| | | | Resolved | Found |
| simPhage | 2 | 0 | 1 | 1 |
| Lake Water | 192 | 8 | 6 | 6 |
| Paddy Soil | 466 | 2 | 7 | 7 |
| Wastewater | 92 | 1 | 1 | 1 |
| IBD | 262 | 2 | 20 | 32 |

### 10. Further benchmarking results

#### 10.1 Bin/genome counts of different quality

Table S8. Benchmarking results of PHAMB (Johansen et al. 2022) and Phables. Best counts are highlighted in bold.

| Dataset | Tool | Number of phage bins/ genomes identified | Number of genomes with < 10% contamination | Number of genomes with > 90% completeness | Number of complete and high-quality genomes | Number of genomes with CheckV warnings |
|---|---|---|---|---|---|---|
| Lake Water | PHAMB | 203 | 203 | 162 | 162 | 23 |
| | Phables | **210** | **209** | **165** | **165** | **1** |
| Paddy Soil | PHAMB | 189 | 189 | 154 | 154 | 96 |
| | Phables | **479** | **479** | **389** | **389** | **3** |
| Wastewater | PHAMB | 62 | 61 | 40 | 40 | 13 |
| | Phables | **94** | **94** | **55** | **55** | **0** |
| IBD | PHAMB | 246 | 239 | 193 | 193 | 75 |
| | Phables | **307** | **305** | **205** | **205** | **5** |
| Lake Water - metaSPAdes | PHAMB | 168 | 165 | 82 | 82 | 17 |
| | Phables | **230** | **224** | **182** | **182** | **0** |
| Lake Water - MEGAHIT | PHAMB | **184** | **183** | 98 | 98 | 17 |
| | Phables | **184** | 182 | **156** | **156** | **0** |

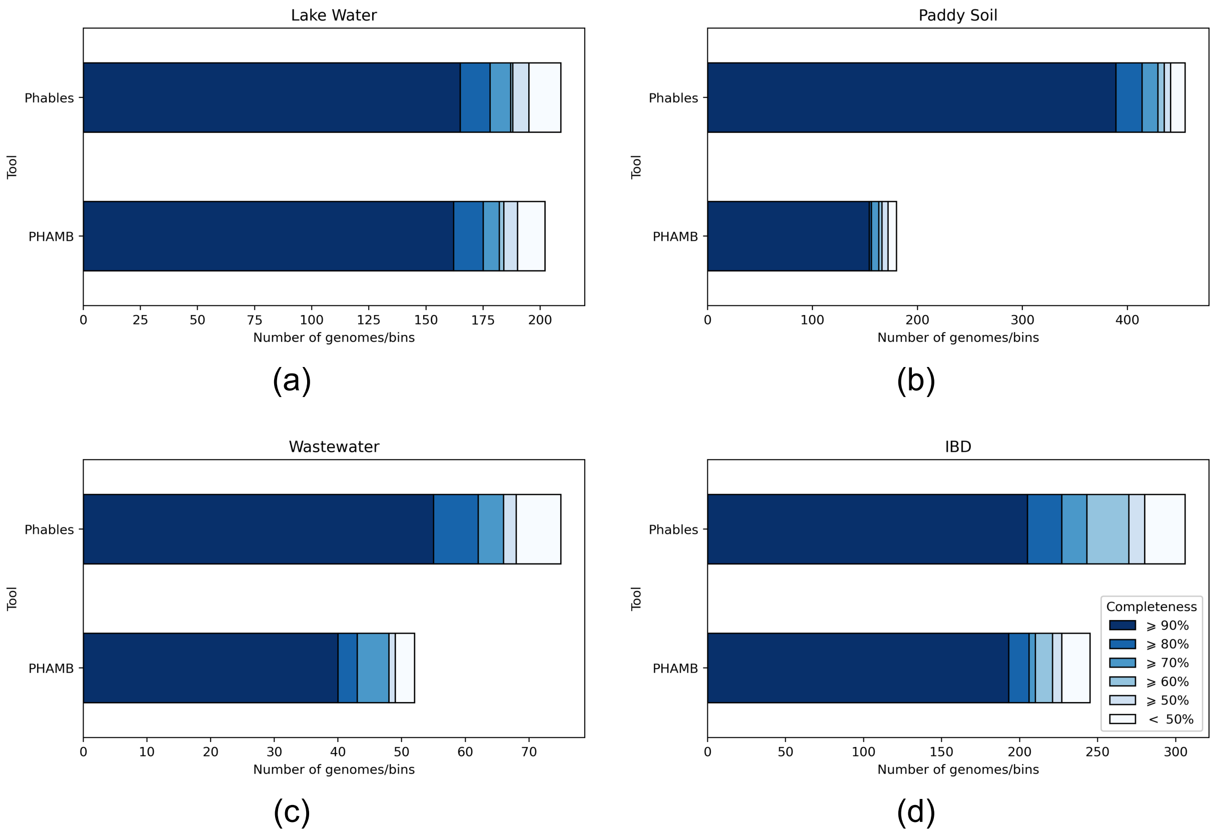


Figure S4: Comparison of CheckV completeness of sequences from Phables and PHAMB (Johansen et al. 2022) for the viral metagenomic datasets (a) Lake Water, (b) Paddy soil, (c) Wastewater, and (d) IBD.

#### 10.2 Quality comparison using all bins containing the selected PHROGs

We benchmarked Phables based on all the PHAMB bins containing the selected PHROGs and the results are presented in Figures S5 - S7. It is clear that even though PHAMB recovers more phage genomes overall, many genomes have CheckV warnings as a result of the erroneous genome structures, high levels of contamination and duplications within genomes due to the presence of multiple closely-related phages. Moreover, it is clear that the majority of the genomes recovered from PHAMB have less than 90% completeness (Figure S7).

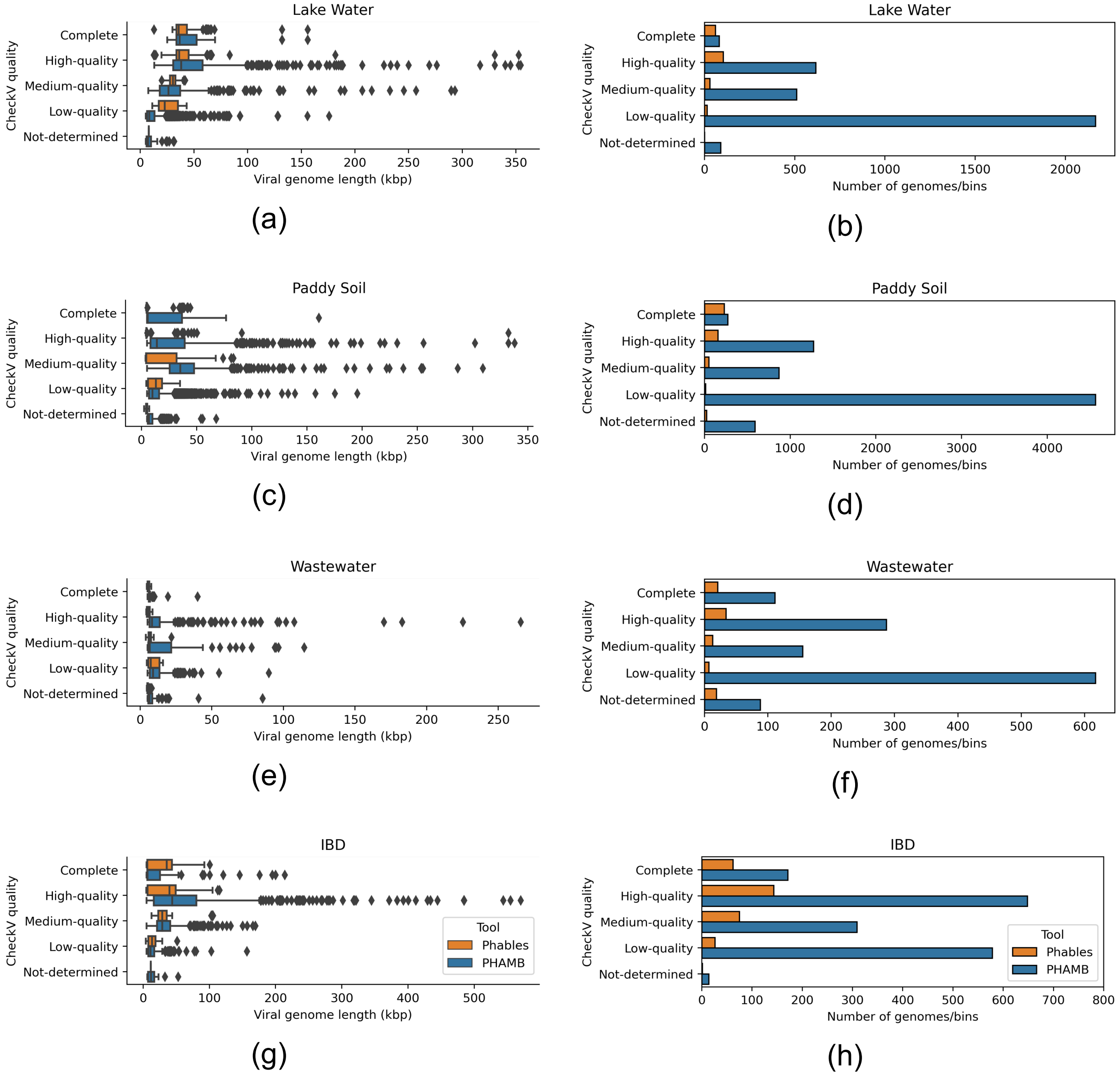


Figure S5: Genome length distribution (first column of figures) and abundance of genomes (second column of figures) belonging to different CheckV quality categories identified by Phables (denoted in orange) and PHAMB (Johansen et al. 2022) (denoted in blue) for the viral metagenomic datasets Lake Water, Paddy soil, Wastewater, and IBD.

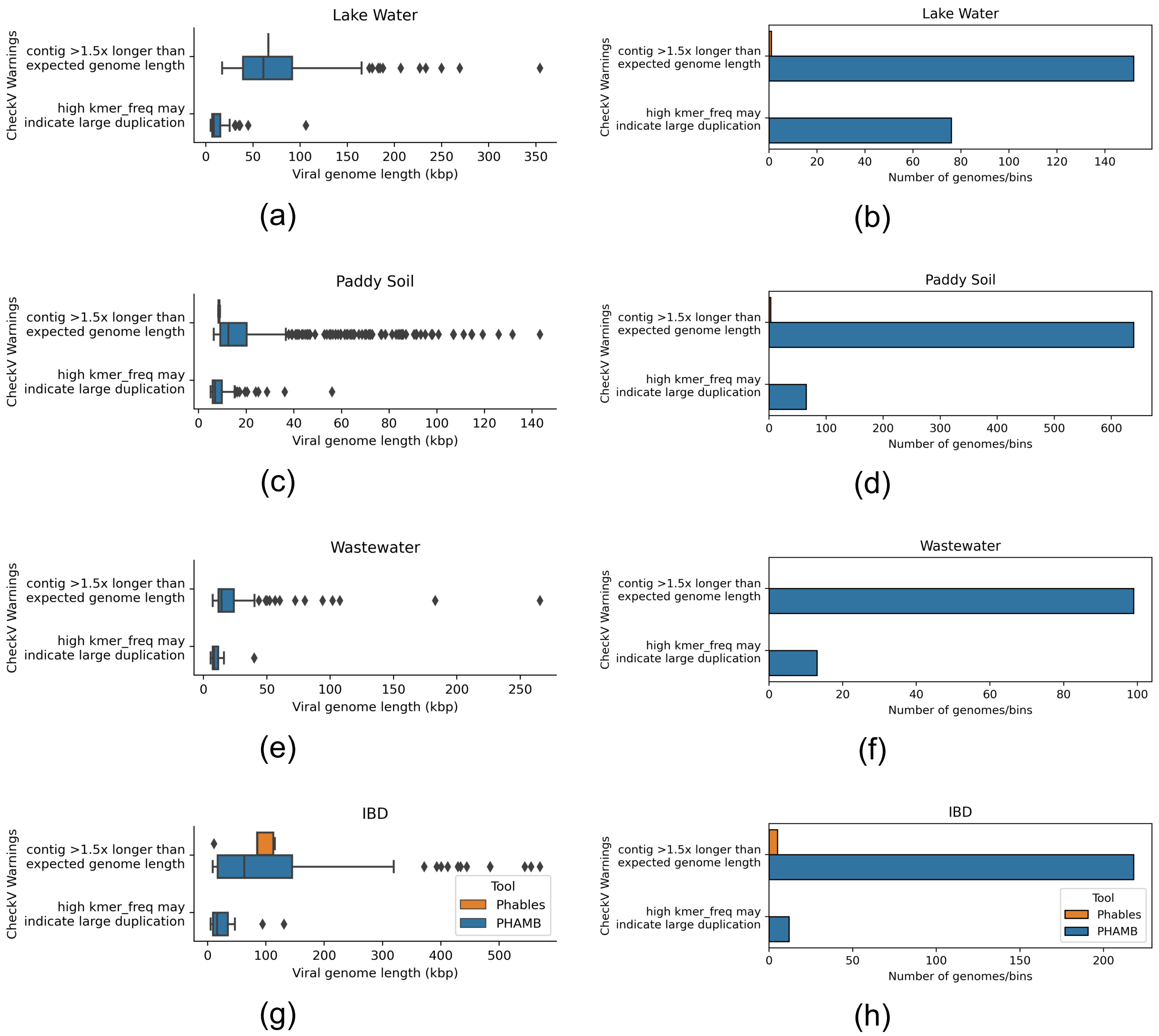


Figure S6: Genome length distribution (first column of figures) and abundance of genomes (second column of figures) having the selected CheckV warnings from Phables (denoted in orange) and PHAMB (Johansen et al. 2022) (denoted in blue) results of the viral metagenomic datasets (a) - (b) Lake Water, (c) - (d) Paddy soil, (e) - (f) Wastewater, and (g) - (h) IBD.

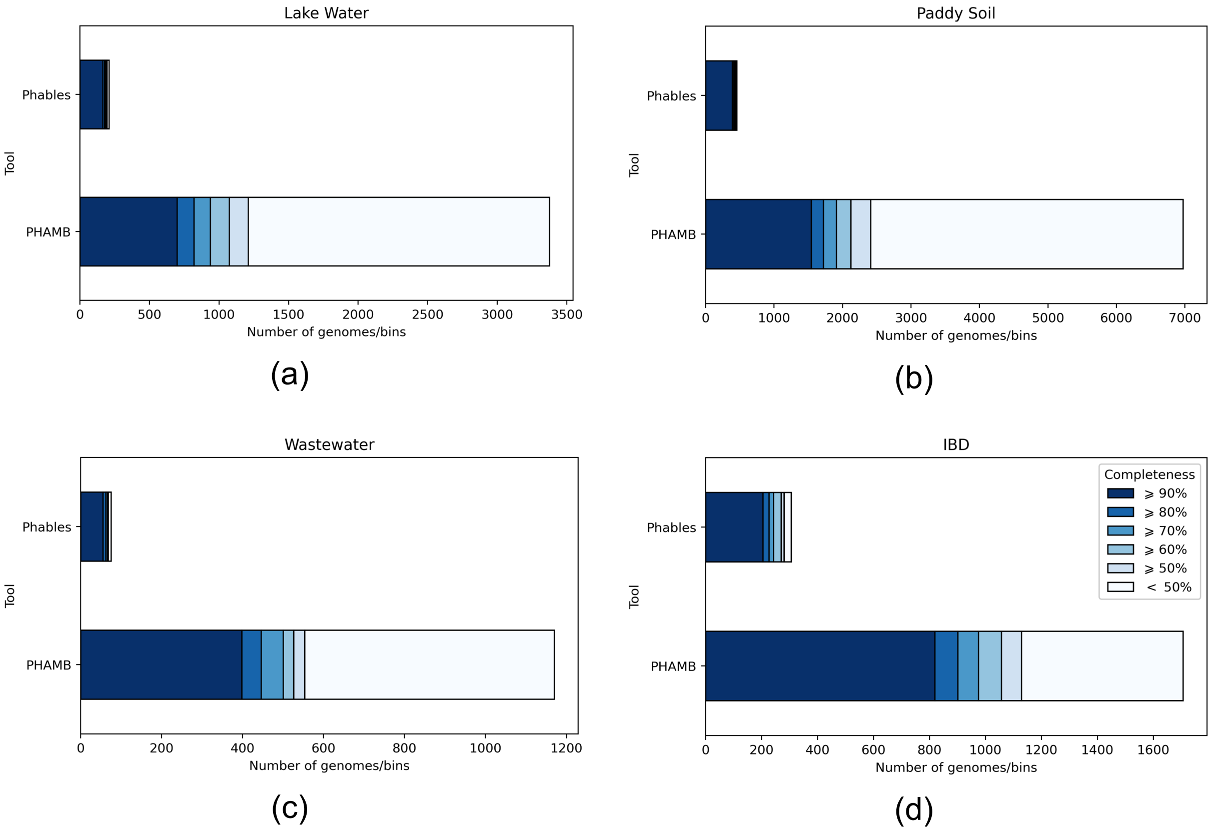


Figure S7: Comparison of CheckV completeness of sequences from Phables and PHAMB (Johansen et al. 2022) for the viral metagenomic datasets (a) Lake Water, (b) Paddy soil, (c) Wastewater, and (d) IBD.

### 11. Complex case 3 phage components

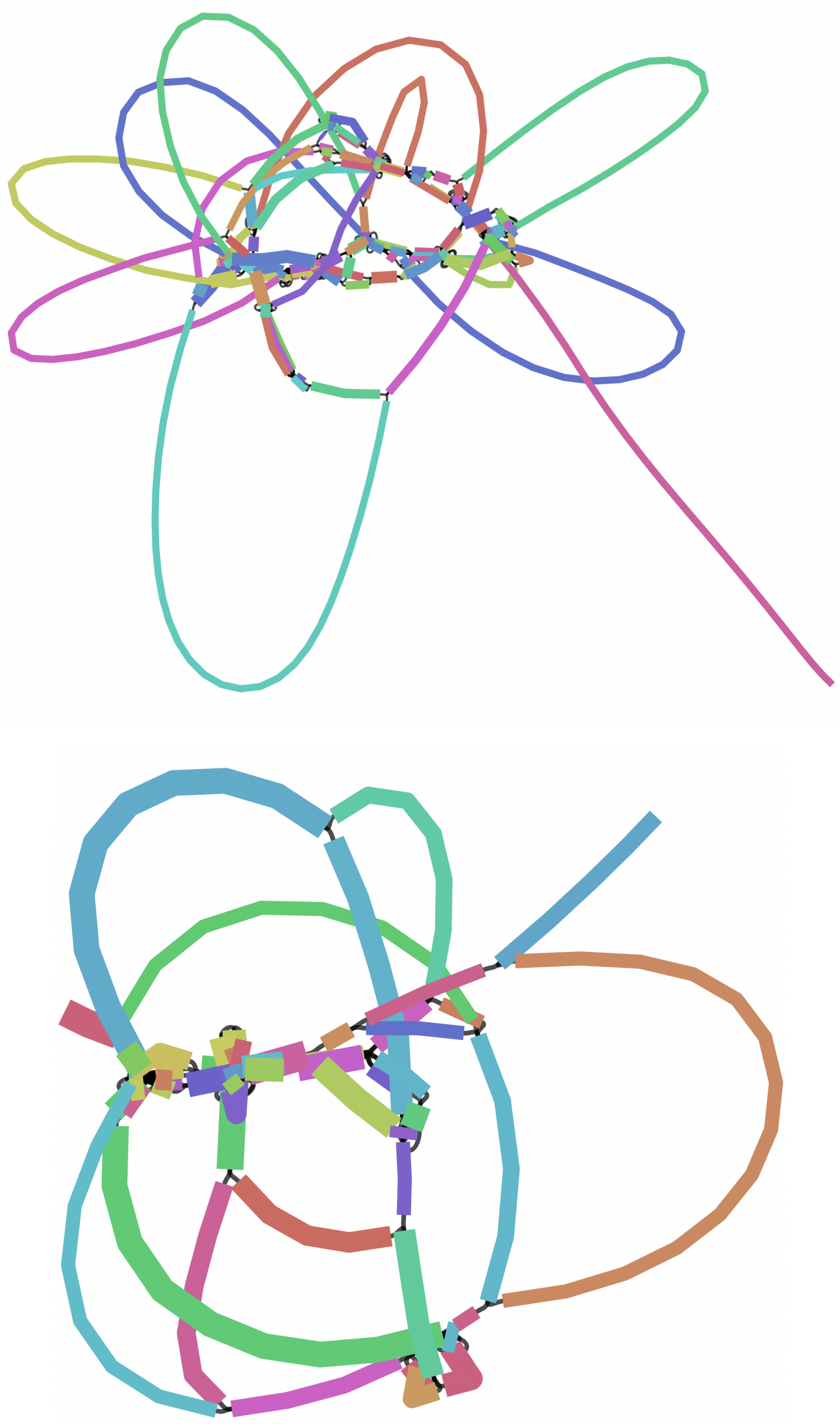

Figure S8. Bandage (Wick et al. 2015) visualisation of two complex case 3 phage components from the IBD dataset where a $st$ vertex could not be found.

### 12. Similarity analysis of resolved genomes from the IBD dataset

We compared the average nucleotide identity (ANI) of 37 resolved genomes from 14 case 3 phage components of the IBD dataset as it had the most number of case 3 components resolved. All pairs have over 95% ANI, and 20 pairs have over 85% alignment coverage.

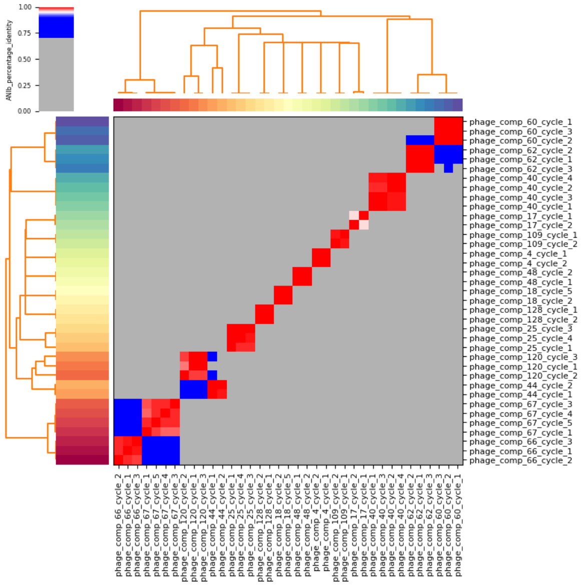

(a)

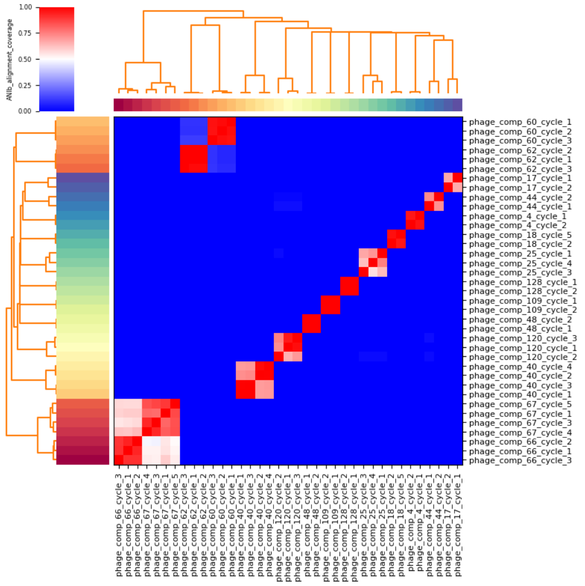

(b)

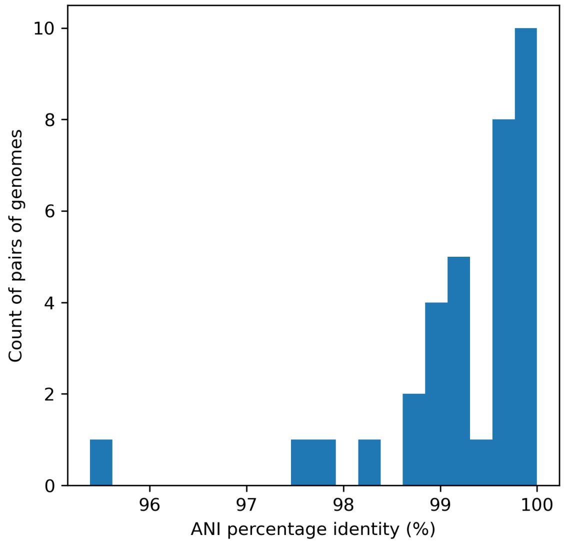


(c)

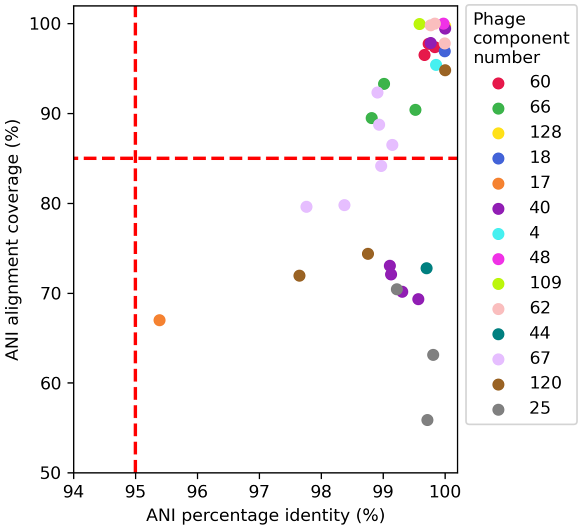


(d)

Figure S9. Average nucleotide identity (ANI) analysis of the pairs of resolved genomes from case 3 components of the IBD dataset. (a) ANI identity percentage, (b) ANI alignment coverage, (c) histogram of ANI values and (d) alignment coverage vs. identity percentage.

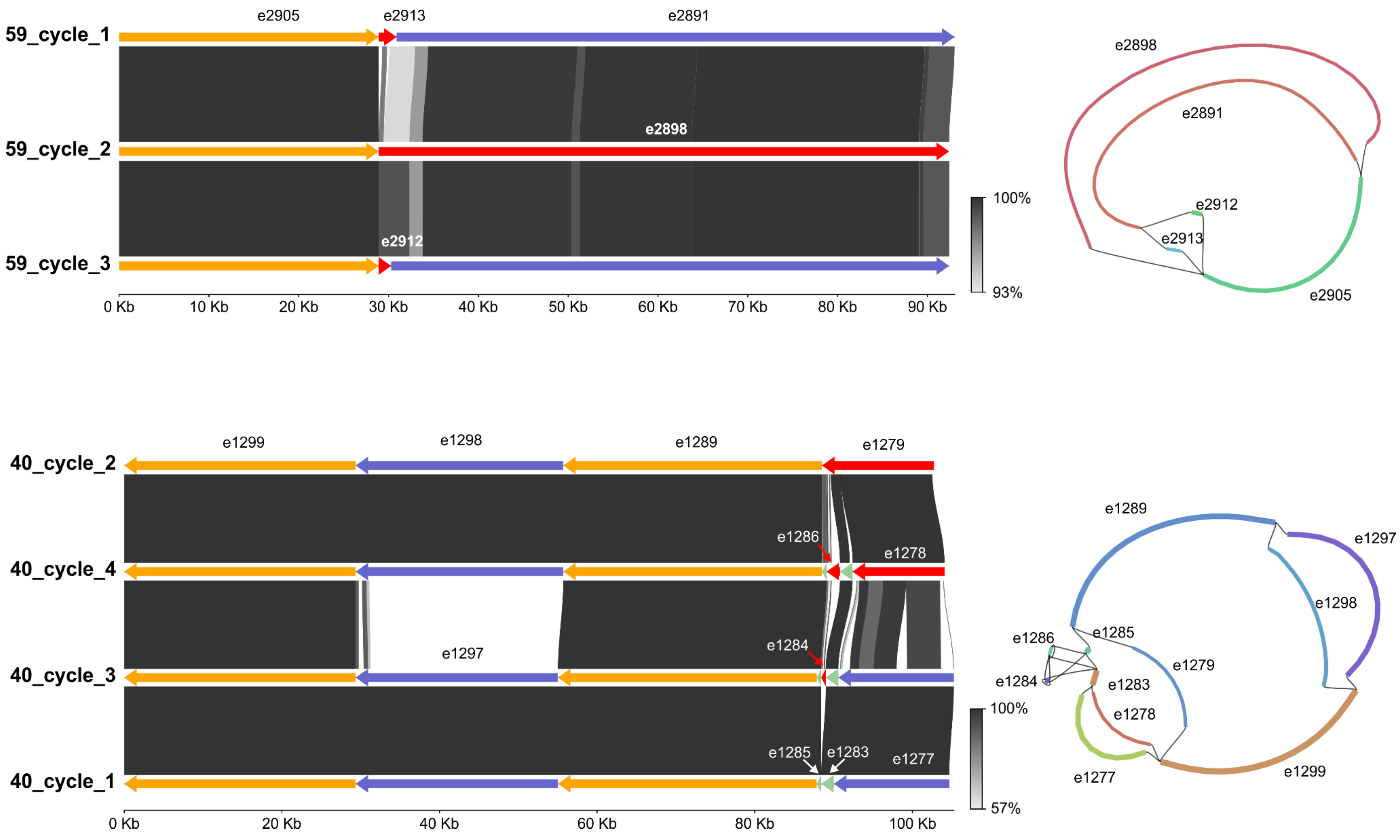


Figure S10: Comparison of similarity between the genomes resolved from two components from the IBD dataset. e[X] denotes the edge numbers from the assembly graph. Red sections denote unique unitigs within the component, blue sections denote unitigs common to two genomes, green sections denote unitigs common to three genomes and orange sections denote unitigs common to all genomes.

### 13. Results from other assembly methods

We assembled the Lake Water dataset using two popular metagenomic assemblers metaSPAdes (Nurk et al. 2017) and MEGAHIT (Li et al. 2015).

Table S9. The number of components found and resolved for different cases of phage components found in the Lake Water dataset assembled using metaSPAdes and MEGAHIT

| Dataset | Case 1 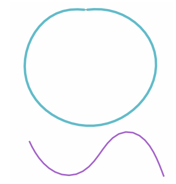 | Case 2 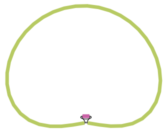 | Cae 3 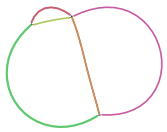 | |
|---|---|---|---|---|
| | | | Resolved | Found |
| metaSPAdes | 34 | 0 | 77 | 105 |
| MEGAHIT | 58 | 0 | 75 | 76 |

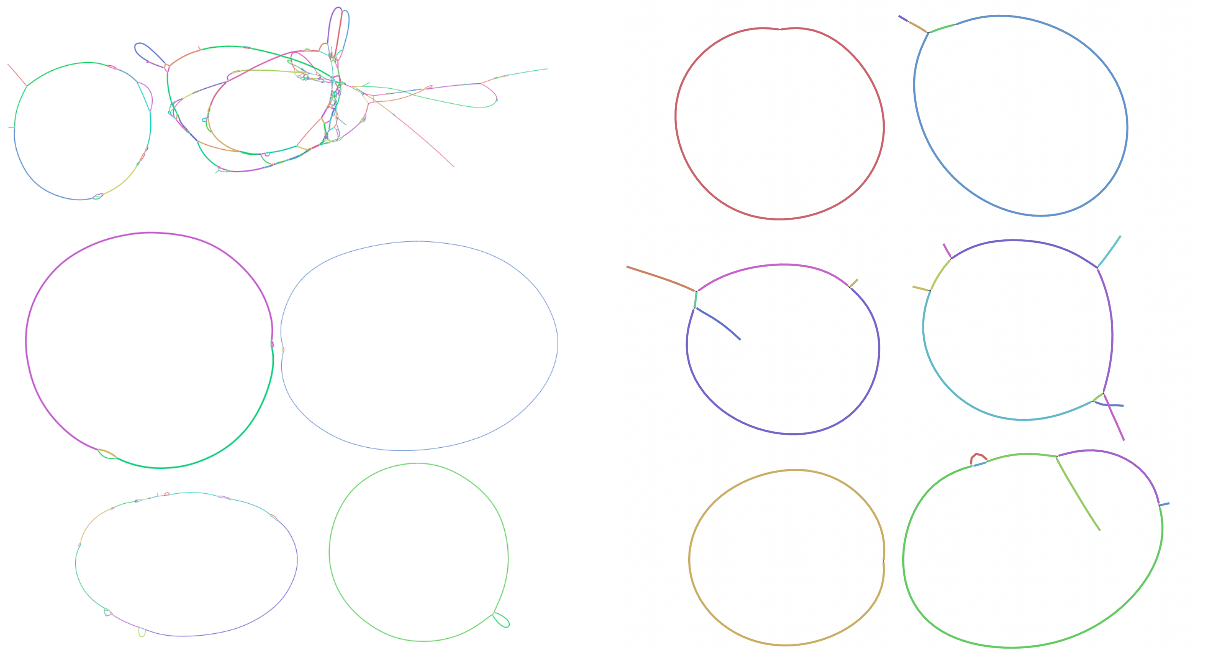

(a) (b)

Figure S11: Bandage visualisations of part of the assembly graph from the Lake Water dataset assembled using (a) metaSPAdes and (b) MEGAHIT.

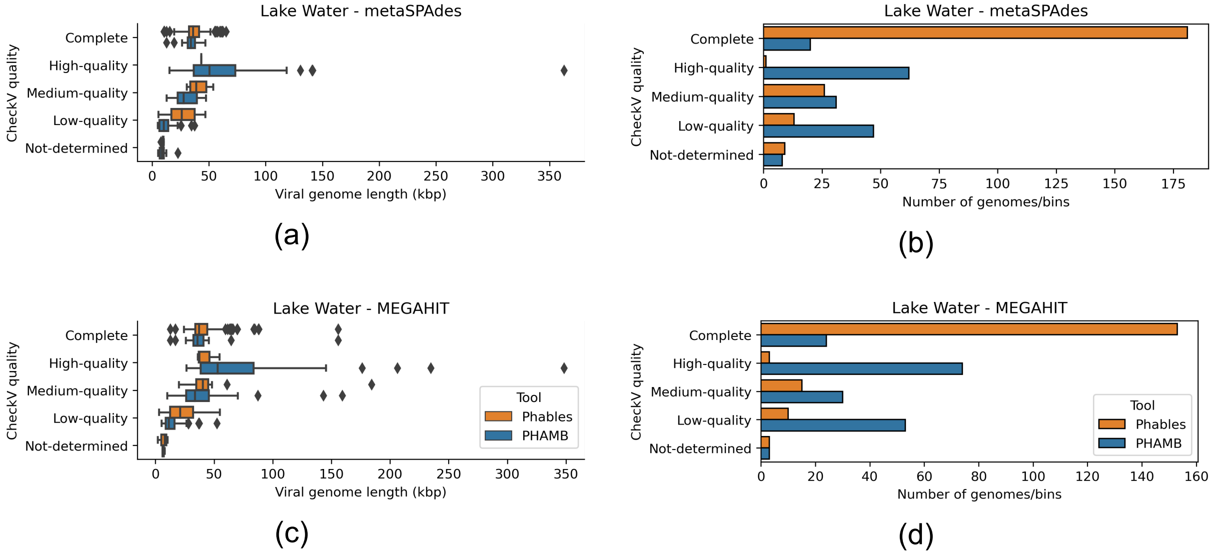


Figure S12: Genome length distribution (first column of figures) and abundance of genomes (second column of figures) belonging to different CheckV quality categories identified by Phables (denoted in orange) and PHAMB (Johansen et al. 2022) (denoted in blue) for the Lake Water dataset assembled using metaSPAdes and MEGAHIT.

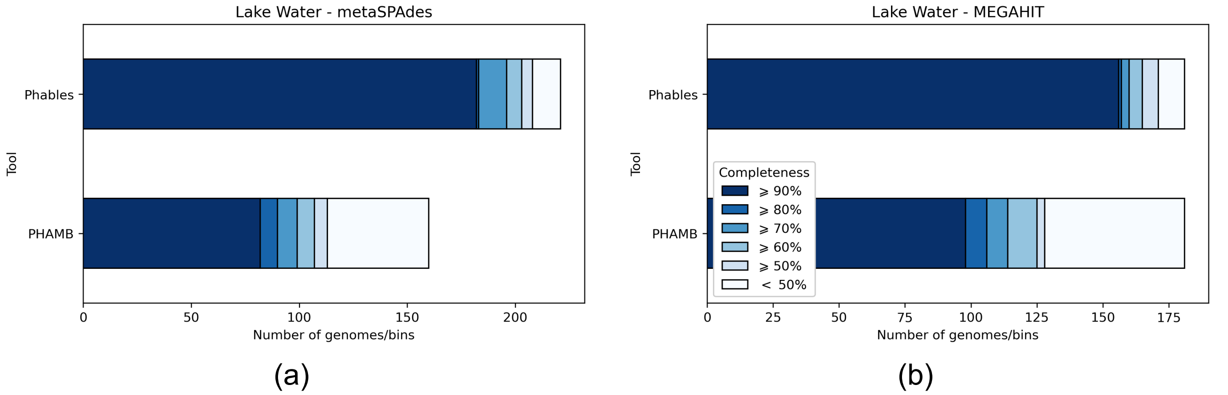


Figure S13: Comparison of CheckV completeness of sequences from Phables and PHAMB (Johansen et al. 2022) results for the Lake Water dataset assembled using (a) metaSPAdes and (b) MEGAHIT.

#### 14. Running time and memory usage

Table S10. Average running time and memory usage of Phables (excluding the preprocessing steps, averaged over 10 runs) with 16 threads for all the datasets. The running time is denoted using minutes (m) and seconds (s), and the memory usage is recorded in megabytes (Mb).

| Dataset | Running time (wall time) | Memory usage |
| --- | --- | --- |
| Lake Water | 9 s | 1298.516 Mb |
| Paddy Soil | 12 s | 973.806 Mb |
| Wastewater | 5 s | 272.720 Mb |
| IBD | 33 s | 455.370 Mb |
| Lake Water - metaSPAdes | 1 m 57 s | 2325.927 Mb |
| Lake Water - MEGAHIT | 1 m 52 s | 3528.125 Mb |

Table S11. Average running time and memory usage of the complete Phables workflow (including the preprocessing steps, except assembly, averaged over 10 runs) with 16 threads for all the datasets. The running time is denoted using hours (h), minutes (m) and seconds (s), and the memory usage is recorded in gigabytes (Gb).

| Dataset | Running time (wall time) | Memory usage |
| --- | --- | --- |
| Lake Water | 1 h 8 m 30 s | 22.989 Gb |
| Paddy Soil | 3 h 10 m 35 s | 43.159 Gb |
| Wastewater | 27 m 18 s | 16.309 Gb |
| IBD | 3 h 30 m 51 s | 16.403 Gb |
| Lake Water - metaSPAdes | 1 h 58 m 7 s | 16.736 Gb |
| Lake Water - MEGAHIT | 2 h 15 m 45 s | 52.981 Gb |

### 15. Analysis of PHROG hits in plasmids

We scanned 34,513 plasmids from PLSDB (https://ccb-microbe.cs.uni-saarland.de/plsdb) (Schmartz et al. 2022) against PHROGs and bacterial single-copy marker genes and found that 25,585 plasmids (74.13%) had bacterial single-copy marker genes. We also found that 8928 plasmids (25.87%) had no bacterial single-copy marker genes but had at least one PHROG hit from the categories: head and packaging, connector, tail and lysis (refer to Figure S15 for plasmid counts for the top 40 bacterial genera). The selection of the four PHROG categories ensures that most phages are recovered (as they are conserved in phages), but some plasmids can be misclassified as phage components.

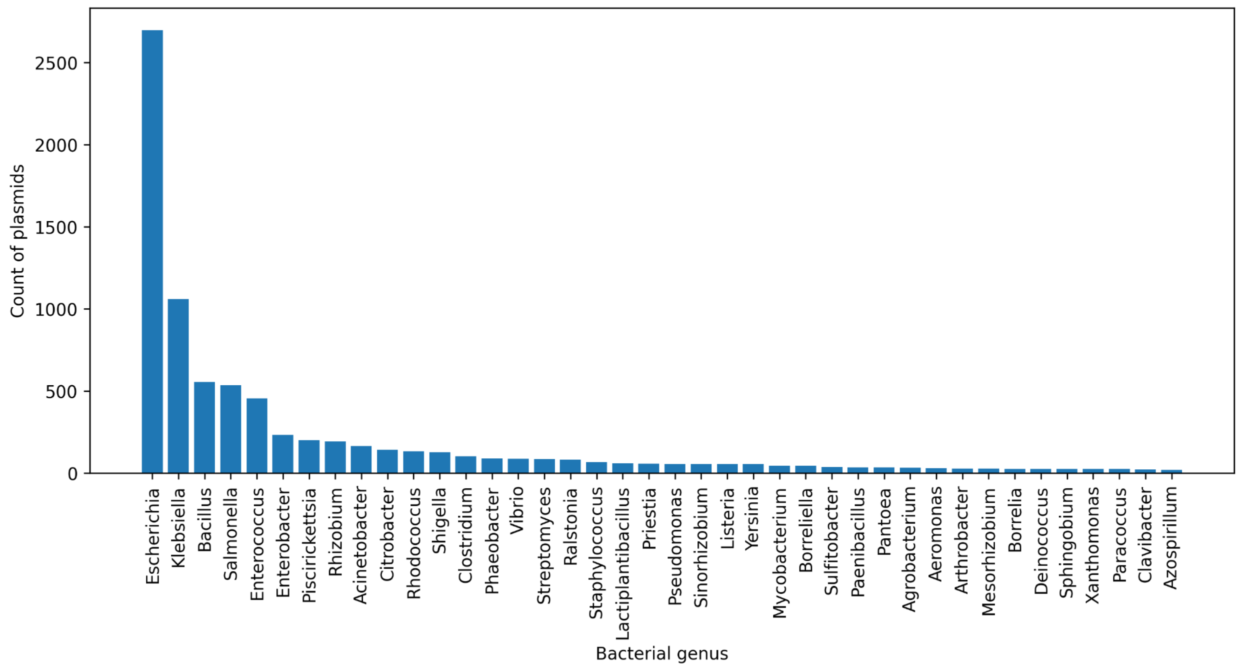


Figure S15: Count of plasmids belonging to the top 40 bacterial genera

Analysis of the selected PHROG categories showed that,

- 705 plasmids (2.04%) have at least one PHROG from all the categories.
- 3793 plasmids (10.99%) have at least one head and packaging PHROG.
- 1052 plasmids (3.05%) have at least one connector PHROG.
- 6988 plasmids (20.25%) have at least one tail PHROG.
- 3393 plasmids (9.83%) have at least one lysis PHROG.

Many temperate phages are found in the host genome as extra-chromosomal plasmids that replicate in line with the cell cycle (Łobocka et al. 2004; Utter et al. 2014; Gilcrease and Casjens 2018), known as *phage-plasmids* (Pfeifer et al. 2021; Pfeifer, Bonnin, and Rocha 2022; Ravin, Svarchevsky, and Dehò 1999), and are elements that are known to be both plasmids and phages (e.g. Escherichia coli bacteriophage P1, Escherichia coli phage N15 or Salmonella phage SSU5). 5363 out of the 8928 plasmids (60.07%) having at least one PHROG hit from the selected PHROG categories are plasmids belonging to bacteria of the genera *Escherichia*, *Enterobacter*, *Salmonella*, *Klebsiella*, *Yersinia*, *Mycobacterium*, *Vibrio*, *Bacillus* and *Clostridiales* (refer to the Figure S16 for plasmid counts of each bacterial genus), which have phages that relate to plasmids encoding proteins homologous to phage sequences (Pfeifer et al. 2021; Gilcrease and Casjens 2018; Dedrick et al. 2016; Octavia, Sara, and Lan 2015; Lan et al. 2009; Myers et al. 2006; Verheust, Fornelos, and Mahillon 2005). Phables can identify such plasmids (or phage-plasmids) as phages and hence, further downstream analysis is required to ensure that the predicted genomes do not include plasmids.

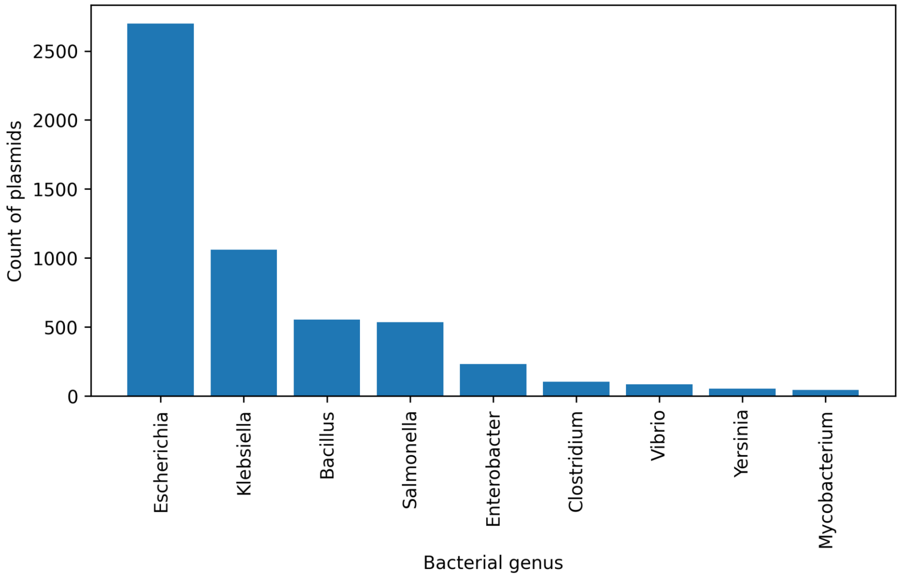


Figure S16: Count of plasmids belonging to bacteria of the genera Escherichia, Salmonella, Klebsiella, Yersinia, Mycobacterium, Vibrio, Bacillus and Clostridiales
